# A transcriptomic atlas of grass senescence reveals divergent underground sink networks limit nitrogen recycling in annuals

**DOI:** 10.64898/2026.05.05.723041

**Authors:** Jonathan O. Ojeda-Rivera, Elad Oren, Sheng-Kai Hsu, Nicholas Lepak, Thuy La, Jingjing Zhai, Michelle C. Stitzer, Abou Yobi, Ruthie Angelovici, Edward S. Buckler, M. Cinta Romay

## Abstract

Senescence enables plants to remobilize and recycle nutrients from aging organs to support growth, reproduction, and survival. In annual crops like maize, nitrogen remobilization from leaves to grain is incomplete, with 30-50% of nitrogen stranded in aboveground tissues and subject to environmental loss. Mitigating nitrogen loss in annual crops could be achieved by leveraging the physiological strategies of perennial grasses, which remobilize nitrogen and other nutrients into underground organs at the end of the growing season, thereby preventing environmental leakage. To uncover the molecular basis of perennial nitrogen recycling to underground organs, we built a transcriptomic atlas from field-grown plants, comprising 2,685 RNA-seq libraries from 14 grass species within the Panicoideae (Poaceae), utilizing maize and sorghum as annual references for comparative analyses. The atlas spans leaves, roots, stalks, and rhizomes across two seasons, from mid-growing season to senescence. Using a photosynthetic index to align the leaf’s transition from nitrogen sink to source across species, co-expression network analysis revealed that the subnetworks driving leaf nitrogen recycling are preserved across annuals and perennials. However, we discovered that the subnetworks associated with underground sink establishment, specifically those associated with seed-like dormancy and desiccation tolerance pathways, have diverged among annual crop accessions. Our work identifies conserved gene candidates and networks that could be used to reintroduce perennial-like nutrient recycling into annual crops to enhance long-term nutrient retention in the field.

## Introduction

Modular growth enables plants to discard and dismantle organs, such as roots and leaves, to fuel the development of new tissues. This dismantling is achieved through senescence, a genetically orchestrated program that mobilizes nutrients out of aging tissues and ultimately leads to their death (Wu et al. 2012). This nutrient recycling is crucial in annual crops, such as maize, where the remobilization of carbohydrates and nitrogen from leaves to ears supports grain yield (Masclaux-Daubresse et al. 2008). However, this remobilization is incomplete, as up to 30-50% of nitrogen remains in aboveground tissues at maturity (Ciampitti and Vyn 2013), where it is subject to environmental loss. Given the scale of maize cultivation, harnessing this residual nitrogen represents an opportunity to circularize nitrogen use and reduce economic and environmental pollution. Understanding the molecular basis of annual and perennial nutrient remobilization is therefore key to engineering strategies to reuse this nitrogen and improve agricultural productivity and sustainability.

The internal processes that control resource allocation in plants have been studied through the lens of sink-source interactions (Sonnewald and Fernie 2018). Source tissues are defined as net exporters of elemental resources that support plant growth, whereas sink tissues are net importers that rely on these inputs for development and function (White et al. 2016). Leaves typically serve as sources of carbon from photosynthesis; however, they also act as nitrogen sinks. In C3 plants, up to 75–80% of leaf nitrogen is contained within the photosynthetic apparatus (Makino and Osmond 1991; Heinemann et al. 2021); this nitrogen investment is reduced in more nitrogen-efficient C4 grasses to approximately 65–75% (Ghannoum et al. 2005, 2010). Leaves transition from nitrogen sinks to sources during senescence, providing an interesting system for studying nitrogen remobilization within plant tissues. In maize, 40-50% of the grain nitrogen is remobilized from leaves and other tissues (Ciampitti and Vyn 2013). Identifying the genes and the spatiotemporal expression patterns that direct this reallocation of nitrogen could provide new avenues for adjusting nitrogen content in grain — either reducing it (Ojeda-Rivera et al. 2025) or increasing it (Maqbool et al. 2021)— depending on downstream grain uses or breeding applications. Tailoring nutrient allocation could increase fertilization precision, reduce inputs, and prevent detrimental environmental impacts from nutrient leakage (Diaz and Rosenberg 2008; Syswerda et al. 2012).

Perennial species possess traits suited for more sustainable nutrient management, characterized by specialized recycling programs that minimize environmental loss and support soil nutrient sequestration (Schwartz and Amasino 2013). Carbon-sink activity is particularly prominent in underground organs of perennial species, which exhibit stronger sink strength than their annual counterparts (Hjertaas et al. 2022). In temperate perennials, such as switchgrass (*Panicum virgatum*), nitrogen is remobilized from above-ground tissues at the end of the growing season and stored in underground organs, including rhizomes and roots (Dohleman et al. 2012). As modified underground stems, rhizomes act as nutrient reservoirs that continue developing beyond grain filling and into dormancy, maintaining a strong and persistent sink that enables temperate perennials to survive stressors such as grazing, drought, or freezing (Yang et al. 2016; Mitros et al. 2020).

In contrast, annual species represent a derived state that evolved from perenniality, prioritizing rapid reproduction under seasonal or terminal stress by shifting resource allocation toward seed production (Friedman 2020). This life history is generally associated with reduced investment in persistent belowground sink tissues, which can limit the capacity for nitrogen recycling relative to perennials. The molecular mechanisms driving the loss or reduction of these recycling traits during the perennial-to-annual transition remain poorly understood in grasses, despite recent progress in defining the genetic basis of perennial nutrient recycling in bioenergy crops (Yang et al. 2016; Mitros et al. 2020).

By around 20 million years ago, declining atmospheric CO_2_ and increasing aridity drove key adaptations in the Andropogoneae, the grass lineage that includes maize and sorghum (Bianconi et al. 2020; Cenozoic CO2 Proxy Integration Project (CenCO2PIP) Consortium*† et al. 2023). These environmental pressures favored the evolution of C_4_ photosynthesis, a highly efficient mechanism for carbon, nitrogen, and water use (Sage and Monson 1998; Sage et al. 2012), and enabled these grasses to expand across warm, open habitats, eventually dominating 17% of global vegetation (Lehmann et al. 2019). The recent sequencing and assembly of multiple Andropogoneae genomes (Stitzer et al. 2025) make it possible to explore the genetic changes underlying their ecological success. In particular, comparative analyses that include temperate perennials such as *Tripsacum dactyloides* — maize’s sister genus (Yan et al. 2019; deWet et al. 1982) — alongside annuals, such as sorghum and maize, offer a powerful framework for investigating the genomic basis of senescence and the regulation of source-sink dynamics.

In this study, we characterize the transcriptional landscape and spatiotemporal dynamics of gene expression in 31 grass accessions across 14 species, from mid-growing season (photosynthesis) to senescence. The dataset primarily features the Andropogoneae tribe, including representatives of *Zea, Sorghum, Tripsacum*, and other ecologically diverse genera, as well as *Panicum virgatum*. Our experimental approach (Figure 1A; see Materials and Methods) captured simultaneous aboveground and underground gene expression patterns in soil-grown plants, yielding a comprehensive resource for systemic analysis of gene expression and nutrient allocation. By utilizing an orthogroup-based approach and a photosynthetic index to align developmental states across species, we utilized crossspecies co-expression networks to identify three conserved programs linked to nitrogen remobilization in perennials: (1) leaf nitrogen recycling, (2) underground structure proliferation, and (3) seed-like dormancy and desiccation tolerance. We provide evidence that while programs 1 and 2 are preserved in annual crops, program 3 was lost in both sorghum and maize through independent annual evolution. Our work identifies conserved gene candidates and regulatory networks that could be leveraged to reintroduce perennial-like nutrient recycling into annual crops, circularize nitrogen use, and enhance long-term nutrient retention in cropping systems.

**Figure 1.**
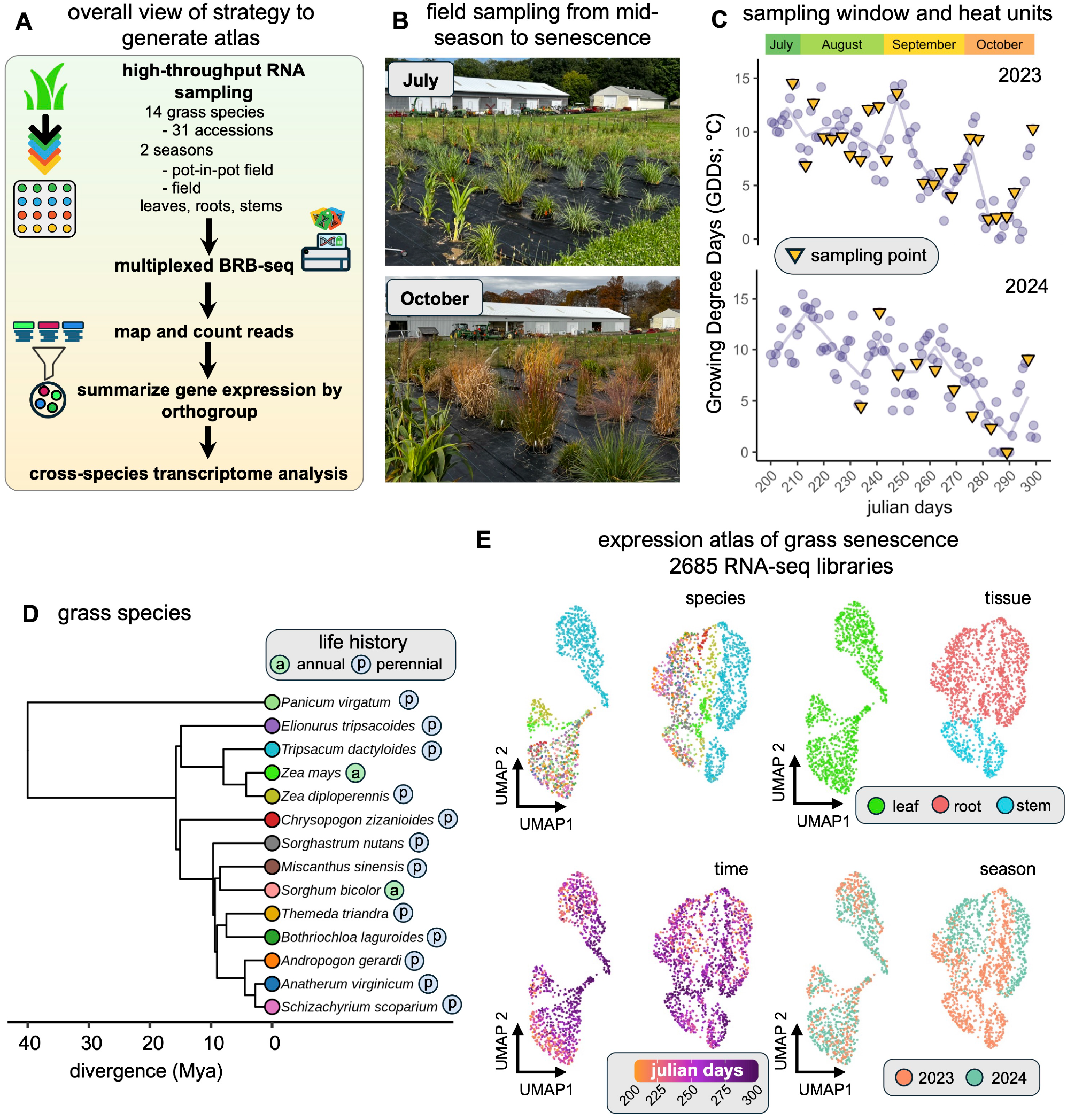
A cross-species, multi-organ, gene expression atlas of grass senescence to explore nutrient recycling. **(A)** Overall view of the multi-species RNA-seq pipeline that was performed to generate the expression atlas. Leaf, root, and stem tissues were collected from 14 grass species (including 31 accessions) across two field growing seasons (years 2023 and 2024). Gene expression was quantified using multiplexed Bulk RNA-Barcoding and sequencing (BRB-seq), summarized by orthogroup, and integrated for cross-species comparative analysis. **(B)** Pictures of plants grown in the field at the middle of the season (July 2023) and late season (October, 2024). **(C)** Daily Growing Degree Days (GDDs; °C) plotted against Julian days for the 2023 and 2024 seasons. Yellow triangles indicate specific sampling events. The variation in GDDs reflects the range of heat units experienced by the field-grown plants, capturing the environmental progression from mid-summer through late-season senescence. **(D)** Phylogenetic tree of the species used in this study. Tips are labeled by life history, annual (a) or perennial (p), with color coding by species. **(E)** Uniform manifold approximation and projection (UMAP) of 2,685 RNA-seq libraries retained after quality control, colored by species, tissue, time, and growing season year. UMAP projections for species are color-coded as in panel D.

## Results and discussion

### A cross-species, multi-organ, transcriptomic atlas for grass senescence and nutrient re - mobilization

To resolve the molecular basis of nitrogen recycling during grass senescence, we first sought to understand how gene expression patterns change in aboveground and underground organs as the plant transitions from photosynthesis to senescence. To this purpose, we generated a comprehensive transcriptomic atlas of 14 grass species during the transition from mid-growing season to senescence using two high-throughput field sampling approaches (Figure 1A-C; see Materials and Methods). The dataset includes major annual crops, such as maize and sorghum, as well as diverse perennial lineages (Figure 1D; Supplementary Dataset SD1). All accessions were soil-grown under field conditions in Ithaca, NY, USA (USDA Hardiness Zone 6a). We sampled leaves, roots, and stalks or rhizomes at 34 sampling points over a 14-week developmental window spanning two growing seasons (Figure 1C).

To ensure consistent gene model quality and minimize annotation bias across diverse genomes, we used deep-learningbased Helixer annotations (Holst et al. 2025) for 13 species, while retaining the *Zea mays* B73 annotation (Hufford et al. 2021) as a functional anchor. Reads were mapped to the respective reference genomes and quantified at the gene level. We generated 3,798 RNA-seq libraries using a high-throughput BRB-seq approach, of which 2,685 were retained after quality control (mapping rate >35% and total mapped reads > 100k). A summary of mapping statistics is included in Supplementary Dataset SD2.

To enable cross-species comparison, we aggregated reads to homologous gene families (orthogroups). Orthogroups were inferred using OrthoFinder (Emms and Kelly 2019) (Supplementary Figure 1). We inferred 97,196 homologous gene families, which captured 90% of annotated genes (Supplementary Figure 1A). To achieve a comprehensive comparative framework, we restricted the orthogroup set to 17,329 families conserved across all tested species (Supplementary Figure 1B). We also needed to define a threshold to minimize the inclusion of genes that are overly duplicated or potentially misannotated. Because many of the sampled species are polyploid, we could not use a simple gene-count cutoff. Therefore, we defined a core set of orthogroups for downstream analysis based on two criteria: (1) the presence of at least one member in all 14 species to ensure evolutionary conservation (Supplementary Figure 1B), and (2) a maximum threshold of 75 gene members per orthogroup (Supplementary Figure 1C). This core set of 16,778 orthogroups captured 72.9% of read counts across the atlas’ libraries (Supplementary Figure 1D). For consistency and to facilitate functional interpretation, the maize gene members of each orthogroup were used as the reference identifiers throughout this work. The orthogroups are available in Supplementary Dataset SD3.

Dimensionality reduction via uniform manifold approximation and projection (UMAP) of the libraries revealed distinct clustering by tissue, species, and developmental time (Figure 1E). A summary of the atlas’s libraries is provided in Supplementary Figure 2A-C. Notably, leaf libraries did not exhibit a significant batch effect between the 2023 and 2024 growing seasons (Figure 1E; Supplementary Figure 2D). In contrast, root libraries showed season-associated clustering, likely due to differences in growing systems between seasons (see Materials and Methods). However, principal component analysis (PCA) of root libraries indicated that this effect was primarily captured by PC4, explaining only 3.64% of the total variance (Supplementary Figure 2E). These results support the dataset’s overall robustness across seasons.

### A cross-species developmental trajectory tracks nitrogen remobilization during grass senescence

From a physiological point of view, the leaf transitions from a nitrogen sink to a nitrogen source during senescence (Wu et al. 2012). Mapping this developmental trajectory (Figure 2A) is key to identifying the genes and regulatory networks associated with nitrogen recycling. However, because multiple exogenous and endogenous factors influence senescence in addition to chronological age (Woo et al. 2019), we sought to identify a common developmental trajectory across species as a reference. For this purpose, we focused our analysis on leaf libraries and modeled the shared transcriptional dynamics across all tested species (Figure 2B) using a pseudotime approach (Campbell and Yau 2018). Pseudotime revealed a shift from the expression of photosynthesis-related orthogroups to senescence across species (Figure 2B). This shift was supported by the decrease in expression of core C4 photosynthesis markers such as *PHOSPHOENOLPYRUVATE CARBOXYLASE* (*PEPC1)* and *RUBISCO SMALL SUBUNIT* (*RBCS1*) (Mendieta et al. 2024) across this trajectory (Figure 2C). In contrast, senescence markers, including *SENESCENCE-ASSOCIATED GENE 12* (*SAG12; Lohman et al. 1994)* and the chlorophyll-degradation regulator *STAYGREEN 1 (SGR1*; Hörtensteiner 2009), were upregulated across this trajectory (Figure 2C).

**Figure 2.**
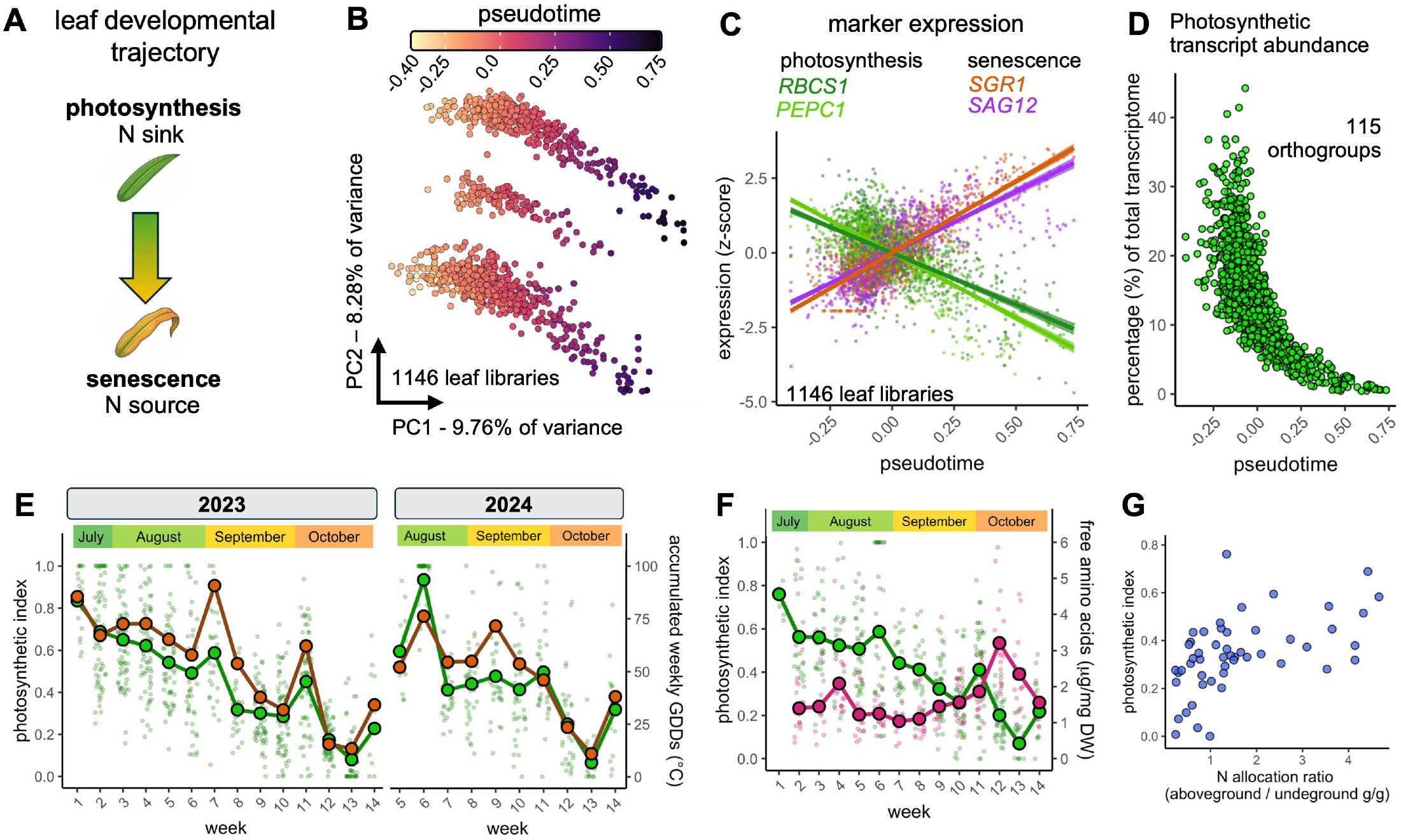
A photosynthetic transcriptome trajectory tracks the nitrogen sink-to-source transition in grass leaves. **(A)** Conceptual model of the leaf developmental trajectory during senescence, illustrating the physiological transition from a nitrogen (N) sink (active photosynthesis) to an N source (remobilization). **(B)** PCA-based pseudotime trajectory of 1,146 leaf libraries across 14 species. Each point represents an individual RNA-seq library, colored by its position along the common developmental trajectory (pseudotime). **(C)** Relative expression (z-score) of photosynthetic (*RBCS1, PEPC1* ; green) and senescence marker genes (*SGR1, SAG12*; purple, orange) across the pseudotime trajectory. Lines represent linear regression fits with 95% confidence intervals. **(D)** Abundance of photosynthetic transcripts as a percentage of the total transcriptome across pseudotime. The 115 orthogroups used for this calculation include core components of the photosynthetic machinery and photosynthesis-related genes (see Materials and Methods). The photosynthetic is derived by scaling this proportion from 1 (maximum photosynthetic abundance) to 0 (minimum). **(E)** Time progression of the photosynthetic index (green; left y-axis) and accumulated weekly Growing Degree Days (GDDs; °C; brown) during the 2023 and 2024 growing seasons. For the photosynthetic index, points represent individual libraries (photosynthetic index; right y-axis), while the trend line shows the mean value for each sampling week. **(F)** Photosynthetic index (green; left y-axis) and total free amino acid concentration (magenta; right y-axis) in *Tripsacum dactyloides* leaves across sampling weeks. The peak in free amino acids coincides with the sharpest decline in the photosynthetic index. **(G)** Photosynthetic index vs nitrogen allocation ratio in perennial species (total aboveground grams (g) of nitrogen divided by total underground grams of nitrogen). The sampled set of species included *T*. *dactyloides, S*. *nutans, A*. *gerardi*, and *P*. *virgatum* (see Materials and Methods for details).

Given that up to 80% of leaf nitrogen is invested in the photosynthetic machinery (Heinemann et al. 2021), we hypothesized that we could track nitrogen sink-source status in the leaf by using the expression of photosynthetic genes as a proxy. To do this across species, we identified 115 photosynthesis machinery and photosynthesis-related orthogroups that are downregulated along pseudotime and estimated their proportion of the total transcriptome (Figure 2C; Supplementary Dataset SD4). Despite comprising less than 1% of the total set, these orthogroups accounted on average for up to 40% of the total transcriptome in actively photosynthesizing leaves (Figure 2D). We scaled this proportion from 1 to 0 to obtain a photosynthetic index (*ps_index)* for all leaf libraries (Figure 2E). We observed that *ps_index* decreased over time and was closely correlated with Growing Degree Days (GDDs; Figure 2E; Pearson’s r = 0.936).

We then decided to explore the relationship between the *ps_index* and nitrogen recycling. Because free amino acids are the primary form in which nitrogen is remobilized out of tissues during senescence (Havé et al. 2017), we analyzed the relationship between the *ps_index* and free amino acids in the leaves of perennial *T. dactyloides* (Figure 2E; Supplementary Figure 3A). We observed a negative association between free amino acids and the *ps_index* (r=-0.2, p-val<0.01; Supplementary Figure 3A), which was particularly pronounced during sampling weeks 10-14 (Figure 2E). A negative relationship between free amino acids and *ps_index* was expected, as senescent leaves have lower *ps_index* scores. Glutamine, a preferred amino acid for nitrogen transport in the phloem sap of plants (Woo et al. 2019), exhibited a negative correlation with ps_index (r=−0.36; Supplementary Figure 3B). Furthermore, the activation of *SAG12* and *SAG15* proteases (Zhang et al. 2014), which release free amino acids for transport, coincided with the free amino acid peak at week 12 and 13 (Supplementary Figure 3C-D).

To further explore the relationship between *ps_index* and nitrogen recycling in perennial species, we determined the change in the aboveground to underground N allocation ratio from September (week 7) to October (week 14; Figure 2F). We observed that the N allocation ratio to aboveground tissues decreases by the end of the growing season (Figure 2F) and that this allocation to aboveground tissues is positively correlated with *ps_index* (Spearman’s ρ=0.57; p-val < 0.01). Lastly, using an independent maize base-to-tip leaf gradient dataset, we found a positive association between the *ps_index* and leaf protein content within the leaf section interval we sampled (Supplementary Figure 4A). We corroborated a negative association (r=-0.75) of the *ps_index* with free amino acids (Supplementary Figure 4B) in this independent dataset. Together, these results establish the *ps_index* as a robust internal control for leaf’s nitrogen status and an indicator of the transition from sink to source that marks the onset of nitrogen remobilization to underground organs in both annual and perennial species.

### Co-expression network analysis reveals or - thogroups associated with leaf nitrogen re - cycling

To identify the gene regulatory networks driving nitrogen recycling, we first categorized accessions by their ability to partition nitrogen to underground organs between sampling weeks 7 and 14 (Figure 3A). As expected, the annual reference *Zea mays* did not increase nitrogen allocation to underground tissues (Figure 3A). In contrast, perennial accessions of *A. gerardi, P. virgatum, S. nutans*, and the ‘Pete’ accession of *T. dactyloides* exhibited a statistically significant increase in nitrogen allocation to underground tissues by the end of the season (Figure 3A). While these end-point measurements confirmed nutrient recycling, they lacked the resolution to capture the progression of the nitrogen remobilization program. To provide this granularity, we leveraged the *ps_index* as a continuous developmental coordinate, which allowed us to map the transcriptional transition from nitrogen sink to source across all 324 leaf ‘remobilizer’ libraries (Figure 3B). Using this framework, we then performed weighted gene co-expression analysis (WGCNA; Langfelder and Horvath 2008) to identify orthogroups that are associated with the leaf’s sink and source states (Figure 3C-F). Orthogroups were clustered into 10 co-expression modules (Figure 3C), summarized by module eigengenes (MEs; Figure 3D-E) that captured the main expression trends across species and developmental trajectory (Figure 3D-E). To visualize the network, we summarized the topological overlap matrix (TOM) features of the orthogroups into a two-dimensional UMAP (Figure 3C). For a complete list of orthogroup module assignments and hub rankings, see Supplementary Dataset SD5. For details on WGCNA implementation and parameters, see the Materials and Methods section.

**Figure 3.**
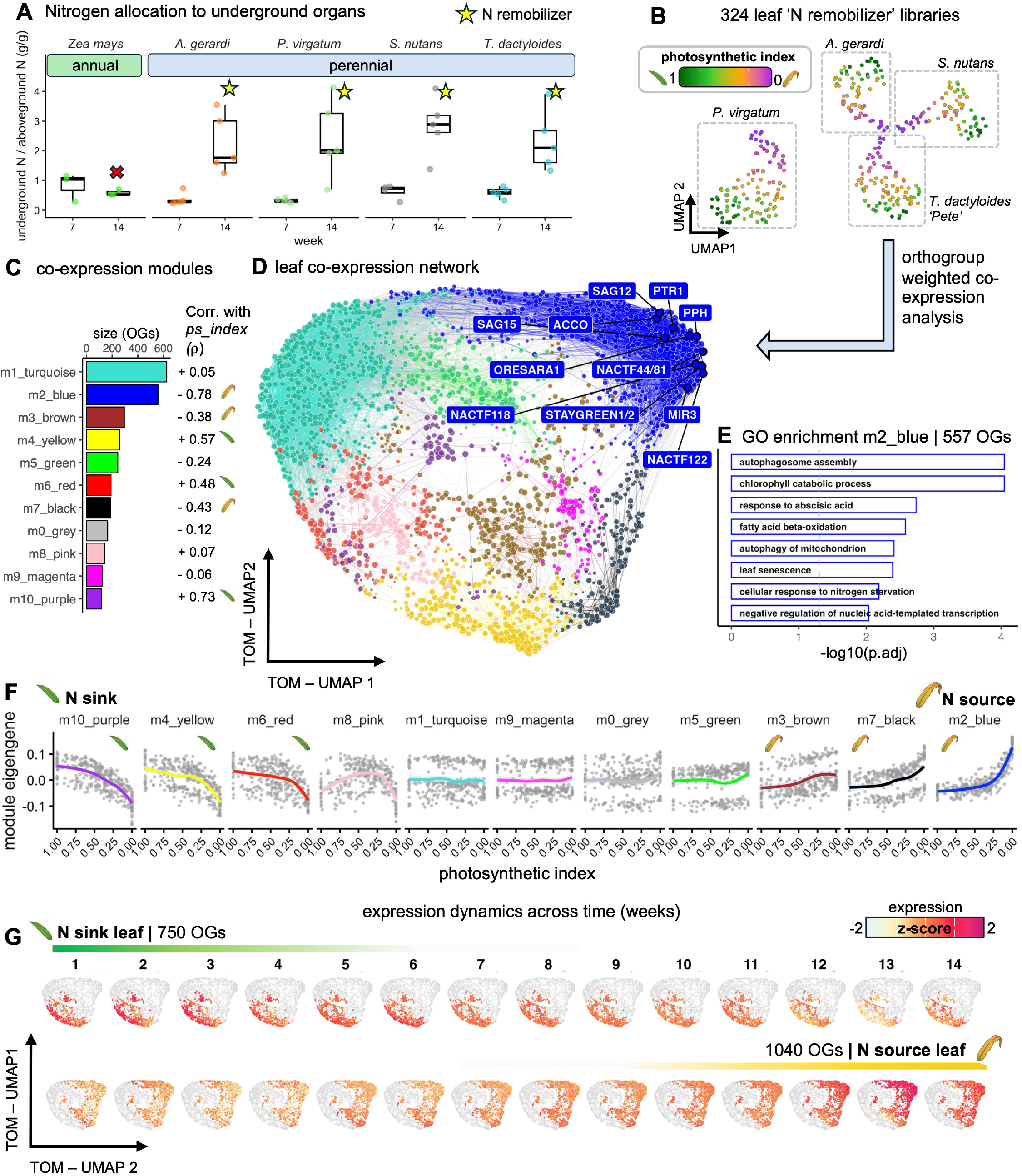
Co-expression network analysis identifies transcriptional programs for leaf nitrogen remobilization. **(A)** Nitrogen (N) allocation ratio (total N in underground tissues / total N aboveground tissues, g/g) measured at sampling weeks 7 (September) and 14 (October). Perennial species identified as “N remobilizers” (stars) show a statistically significant increase in nitrogen sequestration to underground organs (Two-Way ANOVA, Tukey HSD, p < 0.05). In contrast, the annual reference *Z*. *mays* (red x) showed no significant increase (p > 0.05). **(B)** UMAP visualization of 324 leaf libraries from N-remobilize r species, colored by photosynthetic index. **(C)** Summary of the 10 orthogroup (OG) co-expression modules identified by WGCNA. Bars indicate module size (number of orthogroups), and values indicate the Spearman correlation (*ρ*) between the module eigengene and the photosynthetic index (*ps_index*). Leaf icons denote sink-associated (leaf) and source-associated (leaf) modules. **(D)** Topological Overlap Matrix (TOM)-UMAP projection of the leaf co-expression network. Nodes represent orthogroups (OGs) colored by module assignment; key hub orthogroups in the source-associated m2_blue module are highlighted. Maize symbols or synonyms were assigned to orthogroups based on MaizeGDB nomenclature and supporting literature. Highlighted hubs include *ORESARA1, STAYGREEN1/2*, and the following abbreviated OGs: *SAG12/15 (SENESCENCE ASSOCIATED GENE 12/15), PPH (PHEOPHYTINASE), PTR1 (PEPTIDE TRANSPORTER 1), ACCO (1-AMINOCYCLOPROPANE-1-CARBOXYLATE OXIDASE), MIR3 (MAIZE INSECT RESISTANCE 3)*, and NAC transcription factors (NACTF). **(E)** Gene Ontology (GO) enrichment analysis for the m2_blue module (557 OGs). **(F)** Expression trends of module eigengenes (MEs) plotted against the photosynthetic index. Individual points represent libraries; lines represent local polynomial regression fits. Modules are ordered by their correlation to the index, from N-sink (positive correlation) to N-source (negative correlation), as indicated by the green and brown leaf icons. **(G)** Spatial and temporal expression dynamics of sink-associated (750 OGs) and source-associated (1040 OGs) sub-networks (from panel C) across the 14-week growing season. Nodes represent orthogroups. To analyze expression patterns, only the nodes from the N-sink sub-network (top row) or the N-source sub-network (bottom row) are colored; the remaining nodes in the background are shown in grey. Node color indicates the scaled expression (z-score, white to red) within the network topology.

We estimated the association between the eigengene and *ps_index* by calculating the correlation between the eigengene values for each library and its *ps_index* score (Figure 3C). We used these correlation values and expression patterns to classify expression modules as N-sink, N-source, and not associated with N sink-source status (Figure 3F). We found that coexpression network dynamics in chronological time agree with an expected sink-source transition in leaves (Figure 3G). Among the source-associated modules, *m2_blue* (from module 2 ‘blue’) showed the strongest association with the leaf’s N-source status (Spearman’s *ρ* = -0.77, p-val<0.01; Figure 3C). GO analysis revealed *m2_blue* is enriched in categories that include ‘leaf senescence’, ‘autophagosome assembly’, ‘chlorophyll catabolic process’, and ‘transmembrane transporter activity’ (Figure 3E; Supplementary Dataset SD6),suggesting *m2_blue* contains key orthogroups for senescence, autophagy, and nutrient recycling.

Sub-network hubs within *m2_blue* were ranked using module eigengene-based connectivity (kME). The top 25 ranking hubs of *m2_blue* included multiple orthogroups encoding known orchestrators of leaf senescence, such as NAC transcription factors, ethylene, chlorophyll catabolism, and nitrogen remobilization (Havé et al. 2017; Woo et al. 2019), which are highlighted in Figure 3C. For example, we identified multiple NAC transcription factors (Figure 3B; Supplementary Dataset SD5), including the master regulator of age-induced cell death in plants, *ORESARA1* (Kim et al. 2009). We also found an orthogroup containing a 1-aminocyclopropane-1-carboxylate oxidase *(ACCO*)-encoding gene that catalyzes the last step in ethylene synthesis (Houben and Van de Poel 2019), and multiple ethylene response-related transcription factors were identified as *m2_blue* members (Supplementary Dataset SD5). Additional hub orthogroups included those encoding chlorophyll catabolism-related proteins PHEOPHYTINASE (PPH) and SGR1/2 (Woo et al. 2019). Hubs related to N remobilization included *PEPTIDE TRANSPORTER 1 (PTR1)*, which has been linked to the transport of small peptides produced during enzymatic hydrolysis of the storage proteins in the endosperm to the maize embryo (Tnani et al. 2013) and the proteases *SAG12, SAG15, and MAIZE INSECT RESISTANCE 3 (MIR3)*, which have been linked to nitrogen recycling from senescing chloroplasts (Nakabayashi et al. 1999; Zhang et al. 2014) and senescence regulation in maize (Sekhon et al. 2012, 2019). Free amino acid data from leaves show a transient increase in amino acids by the end of the growing season (Supplementary Figure 3A), which coincides with the increase in expression of hub proteases (Supplementary Figure 3C-D). These data highlight this co-expression module as a key set of orthogroups and genes involved in N remobilization from leaves during grass senescence.

To further investigate the molecular components of nitrogen remobilization, we cross-referenced the orthogroups assigned to N sink- and source-associated modules with a curated list of genes involved in nitrogen metabolism and transport (Figure 4). We constructed this reference set by querying MaizeGDB for key enzymes involved in glutamate metabolism (Forde and Lea 2007) and compiling an inventory of nitrogen transporters from the Nitrate Transporter 1/Peptide Transporter Family (NPF; Léran et al. 2014), Ammonium Transporter family (AMT; Dechorgnat et al. 2019), Amino Acid/Auxin Permease family (AAAP; (Lin Deng 2014), and Usually Multiple Acids Move In and Out Transporters (UMAMIT; (Cao et al. 2025). Finally, we identified orthogroups containing cysteine proteases, which are strongly implicated in senescence and nitrogen remobilization (Havé et al. 2017; Wang et al. 2025). This approach allowed us to identify which members of these specialized nitrogen-related pathways are transcriptionally coupled to the sink-source transition. Notably, several nitrogen-related orthogroups were not assigned to either source- or sink-associated modules, indicating they are expressed but not strongly regulated by seasonal source-sink shifts under the conditions tested. The complete list of N-associated orthogroups and maize genes is provided in Supplementary Dataset SD7.

**Figure 4.**
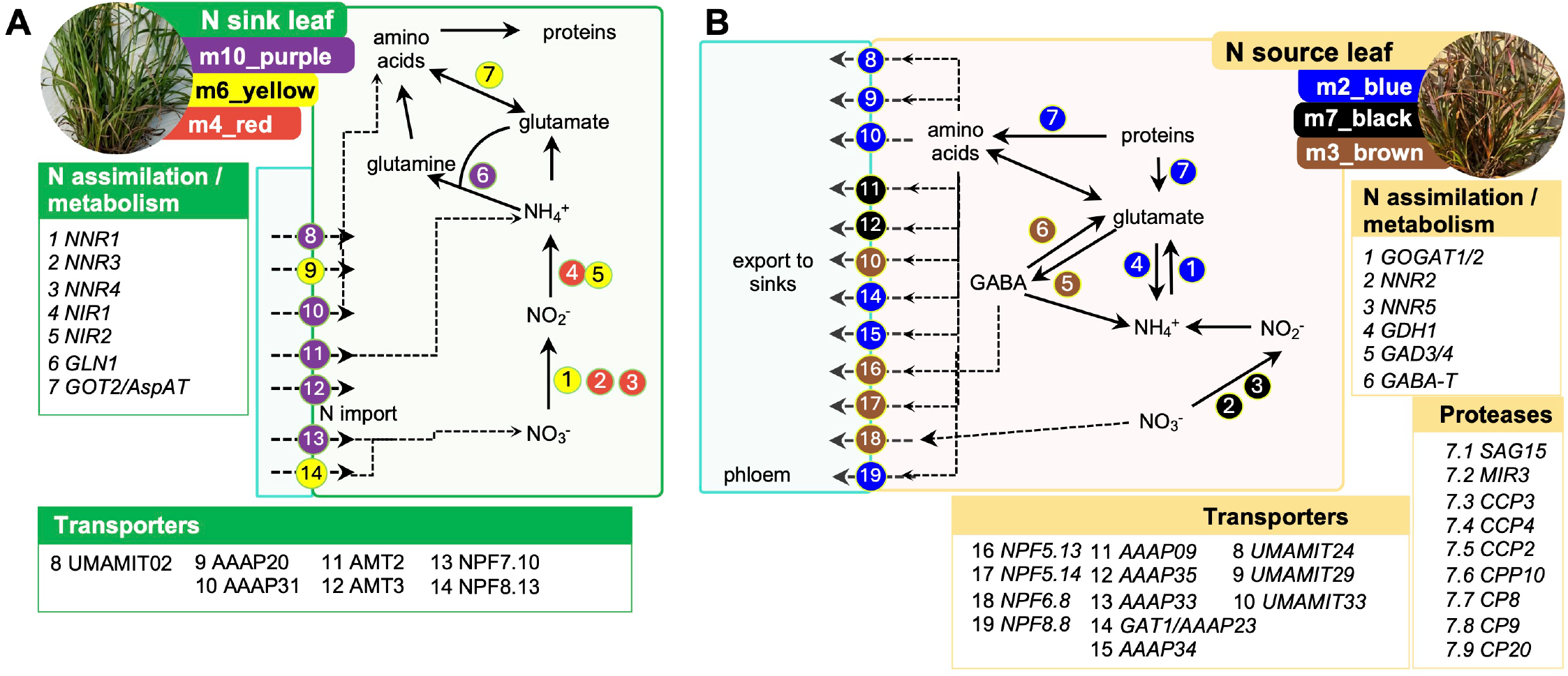
Transcriptional coupling of nitrogen metabolism, transport, and proteolysis to the leaf’s sink-source status. **(A)** Metabolic map of nitrogen assimilation and metabolism in the N-sink leaf state. Numbered circles correspond to orthogroups (OGs) involved in the pathway, with colors indicating their assignment to sink-associated co-expression modules (m10_purple, m6_yellow, m4_red). Processes shown include nitrate reduction, ammonia assimilation, and the synthesis of amino acids into proteins. **(B)** Metabolic map of nitrogen remobilization in the N-source leaf state. OGs are colored by their assignment to source-associated modules (e.g., m2_blue, m7_black, m3_brown). The diagram illustrates protein degradation by senescence-associated proteases, glutamate-GABA metabolism, and the loading of amino acids into transporters (8-19) for export to sinks. Tables below panels A and B list the specific maize gene symbols, proteases, and transporter families assigned to each numbered orthogroup based on literature and MaizeGDB. Enzymes/Metabolism: *NNR (NITRATE REDUCTASE), NIR (NITRITE REDUCTASE), GLN (GLUTAMINE SYNTHETASE), GOGAT (GLUTAMATE SYNTHASE), GDH (GLUTAMATE DEHYDROGENASE), GOT2/AspAT(GLUTAMATE OXALOACETATE TRANSAMINASE / ASPARTATE AMINOTRANSFERASE), GAD(GLUTAMATE DECARBOXYLASE), GABA-T (GAMMA-AMINOBUTYRATE TRANSAMINASE)*. Proteases: *SAG (SENESCENCE ASSOCIATED GENE), MIR (MAIZE INSECT RESISTANCE), CCP/CPP/CP(CYSTEINE PROTEASE)*. Transporters: NPF (Nitrate Transporter 1/Peptide Transporter Family), AMT (Ammonium Transporter family), AAAP(Amino Acid Permease family), and UMAMIT (Usually Multiple Acids Move In and Out Transporters).

### Two distinct subnetworks define proliferative and seed-like nutrient sinks in perennial underground organs

Nitrogen quantification data indicate that underground tissues of ‘remobilizer’ species became nitrogen sinks by the end of the growing season (Figure 3A). To gain insights into the gene regulatory networks underlying the establishment of this late-season nutrient sink that captured N recycled from aboveground organs, we applied WGCNA to underground tissues (roots and rhizomes) using a similar approach to that used for the leaf co-expression network (Figure 5). First, we focused the analysis on ‘remobilizer’ species (Figure 3A). We analyzed a subset of 464 RNA-seq libraries, including root and rhizome libraries from *A. gerardi and S. nutans*, as well as the ‘Pete’ accession of *T. dactyloides*. We couldn’t include *P. virgatum* as most of the root libraries we collected for this species didn’t survive QC (Supplementary Figure 2B; Supplementary Dataset SD2). Second, using the *ps_index* values from the leaves as a reference, we set the corresponding underground libraries along the same trajectory and use *ps_index* as a trait (Figure 5A). Under this framework, underground organs are sinks when the *ps_index* value is close to 0. A UMAP of the 464 RNA-seq libraries shows clustering by tissue species and the *ps_index*-related developmental trajectory (Figure 5A-B). Thirdly, we applied GWCNA to this subset of libraries using *ps_index* as the reference (Figure 5C-E). Orthogroups were grouped into nine co-expression modules (Figure 5C) and named according to their module number and color; however, we added an underground prefix (e.g., *um8_pink*) to avoid confusion with leaf modules. Orthogroup expression was summarized by MEs, which captured the major expression trends across the *ps_index* trajectory (Figure 5D). For a complete list of orthogroup module assignments and hub rankings, see Supplementary Dataset SD8. We identified two main co-expression modules associated with *ps_index, um1_turquoise*, and *um2_blue*, which, interestingly, present opposite expression patterns along the *ps_index* trajectory and time (Figure 5D, H).

**Figure 5.**
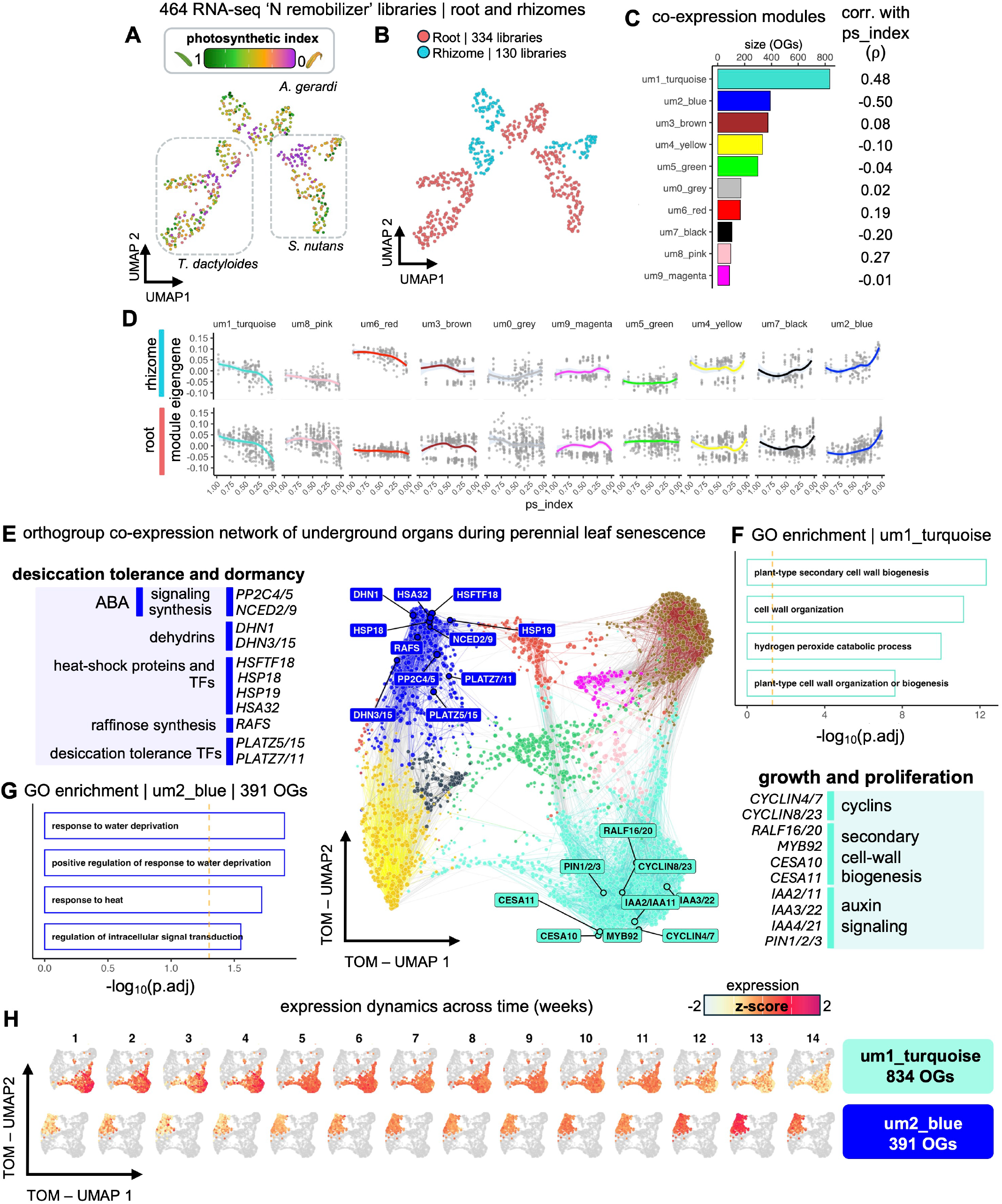
Co-expression network analysis identifies two late-season sub-networks associated with sink establishment in underground tissues. **(A)** UMAP visualization of 464 RNA-seq libraries from perennial underground organs (*A*. *gerardi, S*. *nutans, T*. *dactyloides*), colored by the leaf-derived photosynthetic index (*ps_index)*. **(B)** UMAP of underground libraries colored by tissue type: root (red) and rhizome (blue). **(C)** Summary of nine underground co-expression modules (um0-um9). Bars indicate module size, and values indicate the Spearman correlation (ρ) between module eigengenes and the *ps_index*. OGs (orhtogroups). **(D)** Expression trends of module eigengenes (MEs) plotted across the *ps_index* for rhizome (top) and root (bottom) libraries. Lines represent local polynomial regression fits. **(E)** Orthogroup co-expression network of underground organs. Functionally relevant hubs in the growth-associated *um1_turquoise* and dormancy-associated *um2_blue* modules are highlighted. Acronym definitions for highlighted orthogroups: RALF (RAPID ALKALINIZATION FACTOR), IAA(INDOLE-3-ACETIC ACID responsive protein), PIN (PIN-FORMED auxin efflux carrier), CYCLIN (Cell cycle regulatory protein). Desiccation and Dormancy: ABA (ABSCISIC ACID), PP2C (PROTEIN PHOSPHATASE TYPE 2C), NCED (9-CIS EPOXYCAROTENOID DIOXYGENASE), HSFTF (HEAT STRESS TRANSCRIPTION FACTOR), HSP (HEAT SHOCK PROTEIN), HSA32 (HEAT STRESS ASSOCIATED 32 kDA PROTEIN), DHN(DEHYDRIN), PLATZ (PLANT AT-RICH SEQUENCE-AND ZINC-BINDING PROTEIN), RAFS (RAFFINOSE SYNTHASE), CESA (CELLULOSE SYNTHASE), and MYB92 (MYB-related transcription factor 92). **(F)** Gene Ontology (GO) enrichment analysis for the growth-associated *um1_turquoise* module (834 OGs). **(G)** GO enrichment analysis for the dormancy-associated *um2_blue* module (391 OGs). **(H)** Temporal expression dynamics (weeks 1-14) of the *um1_turquoise* and *um2_blue* sub-networks. Nodes represent orthogroups. Node colors indicate the scaled expression (z-score) across the network topology.

GO enrichment analysis of the *um1_turquoise* module (Figure 5F) revealed enrichment of multiple categories related to secondary cell wall biogenesis (‘plant-type cell wall’, ‘plant-type secondary cell wall biogenesis, ‘cellulose biosynthetic process’, ‘lignin biosynthetic process’) and cell proliferation (‘mitotic cytokinesis’, ‘DNA replication initiation; Supplementary Dataset SD9). Enrichment results suggest roles for this co-expression module in coordinating cell proliferation, elongation, and differentiation in the underground organs of perennial species (Figure 5E-F). Supporting this notion, we identified multiple cyclins that are members of this orthogroup, including *CYCLIN4/7 and CYCLIN8/23*, whose expression is associated with actively dividing tissues and organs in maize (Renaudin et al. 1994; Gautam et al. 2021; Xiao et al. 2021). Top hubs of this module included genes encoding peptides of the *RAPID ALKALIN-IZATION FACTORs (RAFLs; RALF16/20)*, a family of small proteins that have been linked to root development in Arabidopsis (Gonneau et al. 2018) and the regulation of cell wall integrity during pollen tube growth in maize (Zhou et al. 2024). Moreover, we identified an orthogroup that includes *MYB92*. This transcription factor regulates cellulose synthesis and secondary cell wall stability by binding to the promoters of *CELLU-LOSE SYNTHASE 10* and *11*, thereby activating their expression in maize (Zhang et al. 2025); *CESA10/11* are also found in this coexpression module. We also identified multiple auxin signaling-related orthogroups encoding proteins of the IAA, AUX, and PIN families (Supplementary Dataset SD9). From these, orthogroups containing *IAA2/11* and *IAA3/22* are hub orthogroups of this module. Given the predominant role of auxin in coordinating root development (Overvoorde et al. 2010), it is likely that these proteins play roles in the coordination of proliferation, elongation, and differentiation of the underground organs of perennial grasses. Interestingly, enhancing auxin accumulation in maize root tips by overexpressing PIN proteins can improve root growth at the expense of plant height (Li et al. 2018). Overall, GO analysis and functional evidence from maize members of *um1_turquoise* orthogroups suggest this co-expression module orchestrates cell-wall development and proliferation of underground tissues. The expression pattern of the module over time (Figure 5H) suggests that it orchestrates a growth-and-proliferation sink during the mid-growing season, which declines by the end of the growing season as photosynthesis and carbon fixation fall.

Enriched GO categories in the *um2_blue* co-expression module included responses related to tissue dehydration (‘response to water deprivation’, ‘response to heat’), autophagy (SCF-dependent proteasomal ubiquin-dependent catabolic process), and regulatory processes (‘regulation of signal transduction’, ‘negative regulation of cellular macromolecule biosynthetic process). Further exploration of the *um2_blue* orthogroups revealed multiple orthogroups whose members are involved in drought tolerance, seed development, and abscisic acid (ABA) signaling. Top hub orthogroups in this module include a homolog of the *A*.*thaliana HEAT STRESS TRANSCRIP-TION FACTOR C1*, known as *HSFTF18* in maize, which is induced in the maize kernel in response to drought stress (Wang et al. 2019). Other top hubs included members of clade A *PROTEIN PHOSPHATASE TYPE 2C (PP2C4/5)*, which are involved in the ABA signaling pathway (Wang et al. 2018), and 9-cis epoxycarotenoid dioxygenase (*NCED2/9*), which included two gene paralogs of maize’s *VIVIPAROUS14* (Figure 5E). NCEDs catalyze the first committed step in ABA synthesis (Frey et al. 2012) and were first discovered in maize (Tan et al. 1997). Mutants in these enzymes exhibit viviparous phenotypes, characterized by a lack of seed dormancy and higher water-loss rates in detached leaves (Tan et al. 1997; Frey et al. 2012). Orthogroups encoding transcription factors of the AT-rich sequence- and zinc-binding protein family (PLATZ), which have been implicated in seed desiccation tolerance and can confer desiccation tolerance when ectopically expressed in vegetative tissues (González-Morales et al. 2016), are also found in this co-expression module (Figure 5E). The expression pattern of this module over time (Figure 5H), coupled with functional evidence from the literature, suggests that perennial underground organs are leveraging seed-like desiccation-tolerance strategies at the end of the season, most likely to prepare their tissues for winter dormancy.

Further exploration of the gene members of this late-season-expressed module provided additional evidence supporting this seed-like program. For example, another identified hub was *DEHYDRIN1 (DHN1;* Figure 5E), which belongs to the Late Embryogenesis Abundant (LEA) family of proteins which, as indicated by their name, accumulate during embryogenesis and are associated with the establishment of cellular tolerance to dehydration and the multiple abiotic stress linked to this response, including drought, cold, frost, and salt-stress tolerance (Hundertmark and Hincha 2008; Kosová et al. 2021). In maize, *DHN1* has been linked to the drought-tolerance stress response (Zhang et al. 2020) and kernel dehydration (Zhang et al. 2023). Biochemical insights suggest that *DHN1* may change conformation in response to stress and bind other molecules to stabilize cellular components under stress conditions (Koag et al. 2009). It has been proposed that LEAs could protect cellular components by enabling liquid-liquid phase separation, thereby preventing the irreversible aggregation of cellular proteins under osmotic stress (Cuevas-Velazquez and Dinneny 2018). Additionally, we identified another dehydrin orthogroup in the ‘blue’ module, namely *DHN3/15*, whose maize members, *DHN3* and *DHN15*, have been shown to contribute to drought tolerance (Xie et al. 2025) and cold tolerance when heterologously expressed in Arabidopsis and yeast (Chen et al. 2022). Exploration of the carbon-related orthogroups within this module revealed orthogroups *SUCROSE SYNTHASE2/4 (SUS2/4)* and *STARCH SYNTHASES 1/2 (SS1/2)*, which are expressed during grain filling in maize and are key determinants of sink strength (Zhang et al. 2016; Shen et al. 2022). As expected for carbon sinks, rhizomes and roots showed expression of a variety of genes involved in sucrose transport, utilization, and starch biosynthesis. A complete list of orthogroups involved in carbon transport and metabolism is included in Supplementary Dataset SD10.

### Divergence in underground seed-like sub -network distinguishes perennial and annual grasses

The networks built in Figures 3 and 5 provide a reference for sub-networks and orthogroup members that orchestrate aboveground senescence and nitrogen recycling to underground organs in perennials. Because we also sampled annual accessions of *Zea mays* and *Sorghum bicolor* (Figure 1A), transcriptomic data from these accessions offer the opportunity to investigate divergence in the molecular basis of annual and perennial growth and nitrogen recycling strategies. For this purpose, we performed a module preservation analysis in annual libraries (Langfelder et al. 2011) to assess the conservation of N-recycling-associated networks (Figure 6).

**Figure 6.**
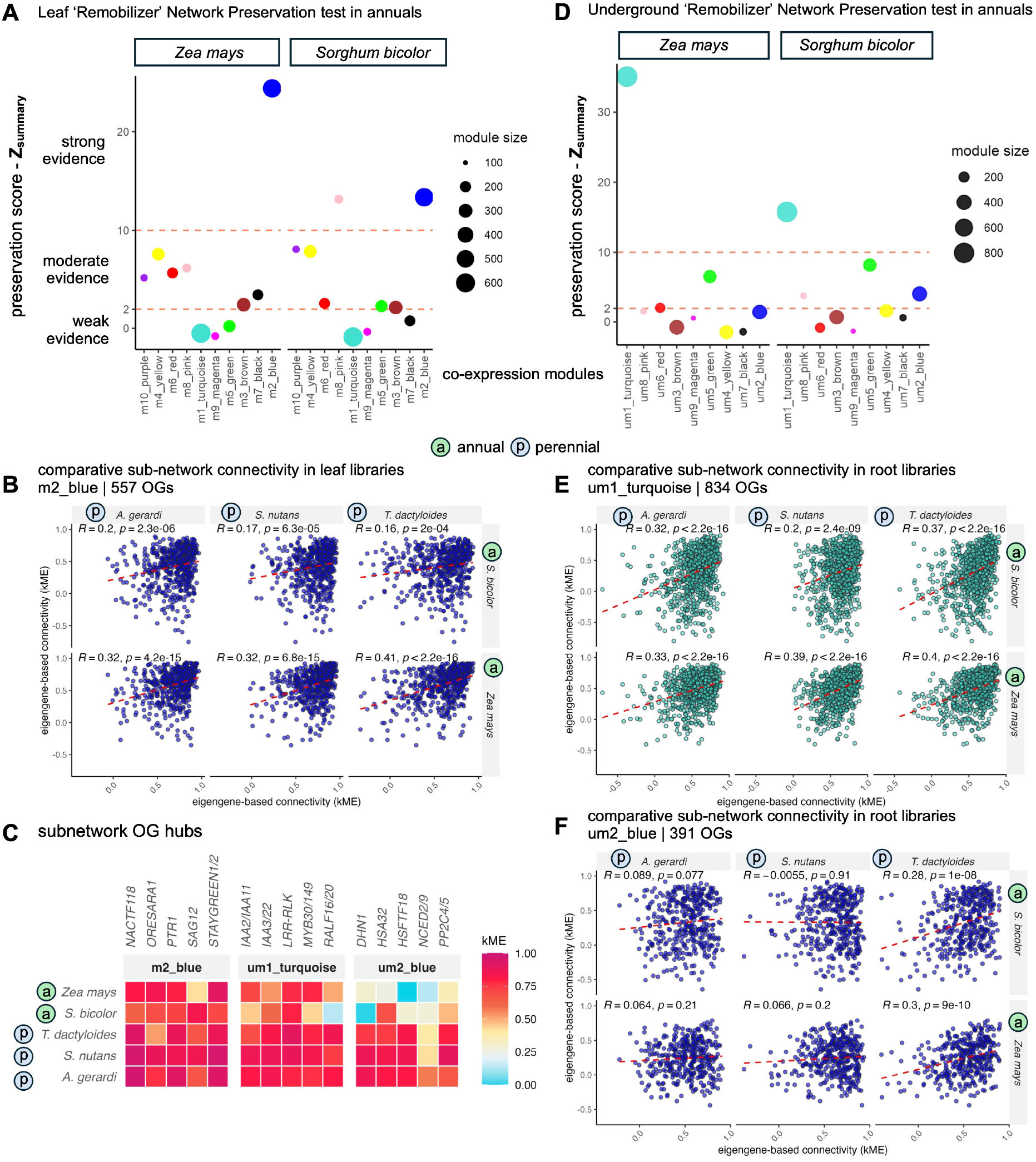
Divergence in underground seed-like sub-network distinguishes perennial and annual grasses. **(A)** Module preservation analysis (Z _summary_) of leaf co-expression modules in annual accessions of *Z*. *mays* (lh244xlh287) and *S*. *bicolor* (BTx623). Dashed lines at Z _summary_ = 2 and 10 indicate thresholds for weak, moderate, and strong evidence of preservation, respectively. The source-associated m2_blue module shows strong evidence of preservation. **(B)** Comparison of orthogroup eigengene-based connectivity (kME) for the m2_blue module between perennial (p; x-axis) and annual (a; y-axis) accessions. Each dot represents an orthogroup that belongs to the subnet. High correlation coefficients and agreement in the upper-right quadrant (high kME in both perennial and annual species) indicate conserved hub status and sub-network structure. **(C)** Heatmap of kME values for representative hub orthogroups across annual (a) and perennial (p) species, demonstrating consistent hub ranking for leaf senescence and underground growth sub-networks, but loss of connectivity for the underground dormancy (um2_blue) sub-network in annuals. **(D)** Module preservation analysis (Z _summary_) of underground co-expression modules in root libraries of annual species. The growth-associated *um1_turquoise* module is preserved, whereas the dormancy-associated *um2_blue* module shows weak evidence of preservation. **(E-F)** Comparison of kME values for the growth-associated um1_turquoise **(E)** and dormancy-associated um2_blue **(F)** modules between perennial and annual species. Note the loss of correlation and agreement in the upper-right quadrant in panel F, particularly for *Z*. *mays* and *S*. *bicolor*.

Regarding the leaf co-expression network, we detected strong evidence of preservation of the *m2_blue* module across annual accessions of *Zea mays* and *Sorghum bicolor* (Figure 6A) based on the summary of network preservation statistics (Langfelder et al. 2011). This co-expression module was the most associated with the leaf’s nitrogen source status (Figure 3C-D, Figure 4B). This result was expected, as annual grasses remobilize nitrogen from senescing leaves, but seeds serve as the primary sink rather than underground organs (Ciampitti and Vyn 2013; Hjertaas et al. 2022). One metric used to assess network preservation is module eigengene–based connectivity (kME), which quantifies the correlation between individual gene expression profiles and module eigengenes (Langfelder et al., 2011). We compared the kME of orthogroups in the *m2_module* between annual and perennial accessions (Figure 6B). The expectation is that if orthogroup connectivity is preserved, it should have a similar kME value in both annual and perennial species. Disagreements in this framework indicate re-wiring or loss of the network’s structure. As expected for a preserved module, *m2_blue* shows strong correlations of kME among annual and perennial species (Figure 6B). Top hubs of this sub-network also present high kME values for both annual and perennial accessions (Figure 6C). This evidence suggests that this sub-network is leveraged across the tested phylogeny and life history syndromes to recycle nitrogen from leaves.

Regarding the underground co-expression network, we detected strong evidence of the preservation of the *m1_turquoise* module in the root libraries of *Zea mays* and *Sorghum bicolor* (Figure 6D), suggesting that this growth and proliferation subnetwork is leveraged by different life-history grasses, regardless of the sampled phylogeny, to establish underground sinks. Comparative kME of orthogroups within this module between annual and perennial accessions shows that the network structure is relatively preserved. Top hubs of the *um1_turquoise* are closely correlated with the module eigengene in annuals and perennials; however, some of them, such as *RALF16/20*, show low correlation with the eigengene (Figure 6C), suggesting some degree of rewiring of this sub-network. A table with kME values for the tested accessions is available in Supplementary Dataset SD11.

Interestingly, we did not find evidence for preservation of the *um2_blue* module in the roots of sorghum and maize accessions (Figure 6D). Comparative kME data showed loss of network structure and connectivity for this sub-network (Figure 6E). Additionally, hub OGs do not correlate with the module eigenvalue of this module in *S. bicolor* and *Zea mays* (Figure 6C; Supplementary Dataset SD11). Moreover, the expression pattern of this module is different in maize with reference to perennials *T. dactyloides* and *S. nutans*, particularly by the end of the growing season (Supplementary Figure 5). Together, these results indicate that while the leaf nitrogen remobilization program and the underground proliferation program are preserved in annual grasses, the sub-network associated with seed-like desiccation tolerance and dormancy has diverged.

## Concluding Remarks

In this study, we present a comprehensive, multi-organ, crossspecies transcriptomic atlas that robustly captures the conserved transcriptional progression of senescence across diverse grass lineages (Figure 1). We provide unprecedented insights into the transcriptional programs of grasses during senescence, particularly for soil-grown roots and rhizomes. This dataset, accessible via our web application (https://maize-genetics.github.io/atlas-of-grass-senescence-dashboard/), serves as a foundational resource for the plant science community to investigate hypotheses related to life history, senescence, and nutrient remobilization in grasses.

By leveraging the dominance of photosynthetic proteins in the leaf proteome (Heinemann et al. 2021) and conserved expression patterns over ∼40 million years of evolution (Figure 2B), we developed a robust photosynthetic index, *ps_index*, to define leaf nitrogen sink and source states (Figure 2). We used this index as a developmental coordinate for network analyses, demonstrating its utility as a standardized proxy for nitrogen retention in leaves and the onset of remobilization across species (Figures 3 and 5). We also demonstrate its transferability for the analysis of other datasets and developmental gradients (Supplementary Figure 4).

Our ancestral gene regulatory networks identify key orthogroups and genes that represent valuable targets for enhancing nitrogen-use efficiency (Figures 3–6). By dissecting sink- and source-associated genes in perennial leaves, we pinpoint known and novel nitrogen metabolism enzymes, proteases, and transporters suitable for biotechnological engineering. These candidates offer specific pathways to either accelerate nutrient remobilization or to support “stay-green” phenotypes in annual crops such as *Sorghum bicolor* and *Zea mays*.

Finally, we characterize the molecular components associated with the establishment of late-season sinks in the underground organs of temperate perennials. Our findings reveal two main programs are required: tissue proliferation and a lateseason seed-like desiccation tolerance (Figure 5). The latter is likely essential for overwintering in temperate environments, as key members, such as LEAs, have been recently linked to freezing tolerance in rhizomes (Oren et al. 2025).

Network preservation analyses showed that while leaf nitrogen recycling and underground growth programs are largely conserved in annual grasses, the desiccation tolerance subnetwork appears largely rewired (Figure 6). Whether this rewiring can be termed a “loss” remains a subject of debate, as it likely resulted in improved fitness for annual species; however, this evolutionary divergence represents the opportunity to engineer perennial-like nutrient sequestration in annual cropping systems. As we have recently proposed, such strategies could keep nitrogen on the farm and reduce environmental loss (Ojeda-Rivera et al. 2025).

Perennial species possess adaptations suitable for sustainable nutrient management, characterized by specialized recycling programs that minimize environmental loss and support soil nutrient sequestration (Schwartz and Amasino 2013). By identifying the specific orthogroups and gene expression dynamics associated with these perennial traits, our study provides a genetic blueprint to reintroduce nutrient recycling into annual crops. Harnessing these ancestral networks offers genetic insights to increase annual nitrogen-use efficiency further, circularize nutrient use, increase soil nutrient sequestration, and enhance long-term agricultural sustainability.

## Materials and Methods

### Plant material and growing conditions

#### Pot-in-pot growing conditions (2023 season)

Accessions of 14 grass species were established in a pot-inpot system designed to standardize soil temperature and moisture across genotypes, enabling root sampling while preventing rooting outside the specified volume and contamination by foreign roots in the field. *Zea mays* and *Sorghum bicolor* seeds were produced in our institutional nurseries, stored at 4 °C, and directly sown in the field. All other species were represented by vegetative clones, which were either propagated directly from wild grassland habitats or derived from the Cornell botanical gardens or seeds acquired through wild collections, the USDA-ARS National Plant Germplasm System, or other sources (Supplementary Dataset SD1). These materials were clonally maintained under greenhouse conditions as described in Hsu et al. (2025), ensuring developmental uniformity at the time of transplant.

Each individual was planted into a 7-gallon (∼26.5 liters) non-woven fabric grow bag (LA TALUS Round Fabric Pots with handles), filled with a modified Cornell potting mix composed of 36.5% vermiculite, 34.6% perlite, 32.8% peat moss, and 6.1% calcined clay (Turface Athletics™ MVP; PROFILE Products, IL, USA) by volume, amended with 0.44% limestone, 0.58% 10-5-10 slow-release fertilizer, and 0.29% calcium sulfate by weight. Fabric pots were nested inside rigid 7-gallon plastic containers (Econo-Grip Nursery Container; A.M.A. Horticulture Inc.), which were fully buried in field soil to match pot depth. Irrigation was delivered via two 2-gallon-per-hour (∼7.5 liters-per-hour) drip emitters per pot, operated daily for 20 minutes to maintain fieldcapacity moisture levels. A schematic representation of the pot-in-pot system is provided in Supplementary Figure 6. Two individuals per accession were established and sampled at the time points shown in Figure 1B.

### Initial greenhouse propagation and field growing conditions (2024 season)

For the 2024 season, seeds were sown in early April in 50-cell trays (SureRoots, T.O. Plastics) using a peat-based substrate (Cornell Soilless Mix). The Cornell Soilless Mix used for initial propagation consisted of two bales (3.8 cu. ft. each; ∼28.3168 L) of sphagnum peat moss, ∼9 kg each of vermiculite and perlite, 1.8 kg of calcium sulfate, 2.3 kg of lime, 1.8 kg of Jack’s Professional 10-5-10 Media Mix Plus III fertilizer, and 0.11 kg of AquaGro 2000 wetting agent. Greenhouse conditions were maintained with a 14-h photoperiod (06:00 to 20:00 h) with supplemental lighting provided by Heliospectra LED arrays; temper-atures were regulated at 28°C (day) and 22°C (night). Following seedling emergence, plants were fertigated with a 15-5-15 Cal-Mag fertilizer at a nitrogen concentration of 100 ppm. Seedlings in propagation trays were moved outdoors to acclimate before transplanting, except for *Zea mays, Sorghum bicolor*, and *Zea diploperennis*, which were direct-seeded by hand in the field. For each species, approximately 20 individuals were established in 3.05-m (10-ft) rows with 1.52-m (60-in) inter-row spacing. Five ranges, including one row from each accession, were randomized and established (total n∼100 plants per accession). Field plots were irrigated with clear water until establishment. Environmental data (Figure 1C) were obtained from Cornell’s Orchard weather station from the NEWA weather resource (https://newa.cornell.edu/all-weather-data-query), and it’s available in Supplementary Dataset SD12. Daily GDD values in Celsius (oC) were estimated using a modified base-temperature method with a base temperature (Tbase) of 10 °C and an upper threshold (Tupper) of 30 °C. The daily GDD was calculated as: GDD=max(0,2min(Tmax,Tupper)+max(Tmin,Tbase)-Tbase). Daily values were then summed starting from the date of planting/emergence to provide the cumulative GDD shown in Figure 1C. Grass accessions excavated from field plots were washed to expose underground organs and dissected into aboveground and underground tissues for RNA sequencing and/or nitrogen quantification (Supplementary Figure 7). Each row within a range was treated as an independent replicate, yielding a total of 5 replicates per accession, sampled at the time points shown in Figure 1B.

### Tissue Collection and RNA Extraction

Tissue for RNA extraction was harvested into 1.2 mL polypropylene 8-strip tubes (96-well tube clusters; Corning) and immediately flash-frozen in liquid nitrogen. For leaf samples, a 10×20 mm section was collected from the midpoint of the leaf blade (equidistant from tip to base), specifically avoiding the midrib. Root samples consisted of five ∼10 mm segments (Supplementary Figure S7B). For stems, including stalks and rhizomes, 10×10 mm sections were harvested from the center of the organ (Supplementary Figure S7B). Samples were homogenized to a fine powder using a Mini-G mechanical grinder (SPEX SamplePrep). Total RNA was extracted from approximately 50 mg of homogenized tissue using the Direct-zol-96 RNA kit (Zymo Research) following the manufacturer’s protocol, including an on-column DNase I treatment. RNA concentration was determined fluorometrically using the QuantiFluor RNA system (Promega), and structural integrity was assessed via an Agilent 2100 Bioanalyzer (Agilent Technologies). Only samples that met the minimum quality thresholds (RIN ≥ 5; concentration > 20 ng/µL) were used for downstream applications.

### Library Preparation and RNA-sequencing

RNA libraries were synthesized using the Mercurius BRB-seq Library Preparation 384 kit (Alpern et al. 2019; Alithea Genomics). Final library quality and quantity were validated using a Qubit fluorometer (Thermo Fisher Scientific) and an Agilent 2100 Bioanalyzer, respectively. Libraries were sequenced on an Illumina NovaSeq X Plus platform using a 25B flow cell at the Cornell University Institute of Biotechnology (Genomics Core Facility). Sequencing was performed in paired-end mode (2×150 bp), targeting a minimum depth of 2 million reads per sample.

### RNA-seq data processing

#### Mapping and counting

Raw reads were demultiplexed using BRB-seq tools (v1.6; https://github.com/DeplanckeLab/BRB-seqTools). Trimming and adapter removal were performed using Trimmomatic (v0.36 in single-end mode with the following parameters: ILLUMINACLIP:TruSeq3-SE.fa:2:30:10, LEADING:3, TRAILING:3, SLIDINGWINDOW:4:15, and MINLEN:36. Post-trimming quality control was conducted using FastQC (Andrews 2010). Ribosomal RNA (rRNA) sequences were filtered by aligning trimmed reads to a custom rRNA reference database (containing *Arabidopsis thaliana* and *Zea mays* rRNA sequences) retrieved from the SILVA rRNA database (Chuvochina et al. 2026) using Minimap2 (v2.27; (i 2018) with the -ax sr preset. The rRNA file is available at https://github.com/jonathan-or/grass-senescence-atlas. Unaligned (clean) reads were extracted using Samtools (v1.2; D(Danecek et al. 2021) and compressed with pigz (https://github.com/madler/pigz). Clean reads were then aligned to the reference genome using STAR (v2.7.10b; Dobin et al. 2013) with the following options: -- twopassMode Basic, --outSAMtype BAM SortedByCoordinate, and --outSAMstrandField intronMotif options. Gene-level quantification was performed using the featureCounts function from the Rsubread package (Liao et al. 2019) in R (v4.2; Core Team 2021). Parameters for counting included isPairedEnd = FALSE, primaryOnly = TRUE, and strandSpecific = 1, with GTF.featureType set to ‘gene’ (or ‘mRNA’ for *Zea mays*) and GTF.attrType = “ID”. Summary statistics, including raw, clean, and mapped read counts, were generated using Samtools stats and custom bash scripts. Alignment summary statistics are included in Supplementary Table SD2.

Reference assemblies and annotations used for mapping

Miscanthus x giganteus libraries were mapped to the *M. sinesis* genome assembly v7.0 (https://phytozomenext.jgi.doe.gov/info/Msinensis_v7_1; Mitros et al. 2020). For *P. virgatum*, we used the var. WBC HAP1 v1.0 as reference (https://phytozomenext.jgi.doe.gov/info/Pvirgatumvar_WBCHAP1_v1_1).

For *S. bicolor*, libraries were mapped to the btx623 reference assembly (Paterson et al. 2009);“https://www.ncbi.nlm.nih.gov/datasets/genome/GCF_000003195.3/”T)

For these three species, we generated gene annotations using Helixer (v0.3.2) with the land_plant lineage model and default parameters (Holst et al. 2025); https://github.com/usadellab/Helixer). Gene annotation files for *P. virgatum, M. sinensis*, and *S. bicolor* are available at https://github.com/jonathan-or/grass-senescence-atlas. For *Zea mays*, we used the B73 v5 assembly and reference annotations (https://maizegdb.org/genome/assembly/Zm-B73-REFERENCE-NAM-5.0; Hufford et al. 2021). For all other species, we used the genome assemblies described in (Stitzer et al. 2025). Assemblies and Helixer annotations for these genomes are available from the PanAnd repository (https://download.maizegdb.org/Genomes/PanAnd/). *Chrysopogon zizanoides* libraries were mapped and counted using the *Chrysopogon serrulatus* assembly as a reference and annotation from the PanAnd repository.

### Orthogroup inference, core set definition, and count aggregation at the orthogroup level

To establish a comparative framework across the 14 species studied, we performed orthogroup inference using OrthoFinder (v2.5.4; Emms and Kelly 2019). Input protein sequences (.faa) for OrthoFinder were extracted from the corresponding GFF3 and FASTA files using gffread (v0.10.4; options: -y; Pertea and Pertea 2020). For *Zea mays*, a custom Python script was used to filter the GFF3 file (B73 v5) for primary transcripts (defined by the “canonical_transcript=1” attribute), and protein sequences were then extracted from this subsetted GFF3 using the same gffread command. OrthoFinder was executed with parameters -S diamond -M msa -t 36 -a 36. Briefly, all-versus-all sequence comparisons were conducted using DIAMOND (v2.0.15, Buchfink et al. 2021) with an e-value threshold of 1×10^-5^. Orthogroups were clustered using the MCL algorithm with a default inflation parameter of 1.5. Phylogenetic relationships were inferred using the Multiple Sequence Alignment (MSA) workflow within OrthoFinder.

Following orthogroup inference, the Orthogroups.tsv output file from OrthoFinder was filtered to retain a core set of orthogroups present across all 14 species. To minimize noise from highly expanded gene families or potential misannotated genes from Helixer, we further restricted the dataset to orthogroups containing ≤75 total gene members. Within each library, raw counts from featureCounts were aggregated at the orthogroup level by summing the counts of all constituent genes within each bin. This process resulted in a unified expression matrix of core orthogroups, which was used in downstream analyses. The core set of orthogroups is included in the Supplementary Dataset SD3.

### Dimensionality Reduction and UMAP Visualization

To visualize the transcriptional atlases’ clustering, we performed Uniform Manifold Approximation and Projection (UMAP) using a pipeline inspired by the Seurat workflow (Hao et al. 2024). First, the core orthogroup count matrix was normalized using a variance-stabilizing transformation (VST) with DESeq2 (v1.46; Love et al. 2014). To focus the embedding on biologically informative signals, we selected the top 3,000 most variable orthogroups. These features were centered, scaled, and capped at a maximum z-score of 10 to reduce the influence of extreme outliers. Principal Component Analysis (PCA) was performed on the scaled matrix using the prcomp function from the R stats package (R Core Team 2021). The first 6 PCs (selected via elbow plot inspection) were used as input to the UMAP algorithm (McInnes et al. 2018), which was implemented using the uwot R package v0.2.3 (https://cran.rproject.org/package=uwot). The UMAP embedding was generated using a cosine distance metric, with n_neighbors=50 and min_dist=0.4 to balance the preservation of local and global structures. Final visualizations were rendered using ggplot2 (Wickham 2016).

### Functional Annotation and Gene Ontology (GO) Enrichment

#### Functional Annotation of Orthogroups

Functional annotations for orthogroups were developed using a multi-step pipeline. For each species, protein sequences were annotated with Gene Ontology (GO) terms using the PANNZER web server (Törönen and Holm 2022) using default parameters. Raw PANNZER outputs files (GO.out) were processed using custom R scripts to generate species-specific Gene Matrix Transposed (GMT) files for Biological Process (BP), Molecular Function (MF), and Cellular Component (CC). Species-level descriptions of GO terms were aggregated into GMT files using a custom Python script. Orthogroup-level annotations were derived by mapping gene-level GO terms to the core orthogroup set using a gene-to-orthogroup mapping table (Supplementary Dataset SD3). Only distinct GO-Orthogroup pairs were retained to prevent redundancy. Output.gmt BP, MF, CC, and GO term description files were used to test enrichment. Orthogroup annotation files are available at https://github.com/jonathan-or/grass-senescence-atlas.

#### GO Enrichment Analysis

Gene Ontology enrichment analysis was performed using the clusterProfiler R package (v4.14.4; Xu et al. 2024). Enrichment was assessed using a hypergeometric test via the enricher function, with a background set comprising all orthogroups in the core expression matrix that have GO annotations. P-values were adjusted (P.adj) for multiple testing using the Benjamini-Hochberg (BH) method. Terms were considered significantly enriched at P.adj < 0.05.

### Pseudotime Analysis

To model the dynamics of the transition from photosynthesis to senescence across species, we performed pseudotime analysis using the PhenoPath R package (v1.30; Campbell and Yau 2018) on leaf-tissue libraries (Supplementary Figure 8). Initial PCA of leaf libraries revealed clustering primarily by species (Supplementary Figure 8A) and chronological time (Supplemental Figure 8B). Notably, the principal component associated with time explained the largest proportion of total variance (PC1; 9.76%), indicating that time-dependent transcriptional changes represent the dominant biological signal in the dataset (Supplementary Figure 8B).

Orthogroups were filtered to retain those expressed in at least 10% of libraries, and normalized to vst-counts as described above. A common developmental trajectory was inferred by fitting a Bayesian linear model where the transcriptional state was modeled as a function of a latent pseudotime component and species as covariate (Supplementary Figure 8C; z_init = 1; ELBO tolerance = 1 × 10^-5). The resulting pseudotime values correlated strongly with chronological time (Pearson’s r = 0.65; Supplementary Figure 8D). To characterize the biological signals captured by the trajectory, we examined the association between orthogroup expression and pseudotime (m_lambda parameter; Supplementary Figure 8E). The top 1% of orthogroups with the strongest positive or negative associations with pseudotime were identified, and subsequent GO enrichment analysis provided biological validation of the trajectory (Supplemental Figure 8F-G). Specifically, chloroplast- and photosynthesis-related GO categories were significantly enriched in orthogroups downregulated along the pseudotime trajectory (Supplemental Figure 8F). The top 1% of cross-species pseudotime-modulated orthogroups are included in Supplementary Dataset SD13.

### Photosynthesis Index Calculation

To quantify the physiological progression of the leaf from N-sink to N-source, we developed a photosynthetic index derived from the core set of photosynthesis-associated orthogroups (Supplementary Dataset SD4) identified in the pseudotime analysis (described above) as being in the bottom 1% of pseudotime effects (Supplementary Figure 8E). This 217-orthogroup set was further screened for associations with photosynthesis-related GO terms. A total of 43 GO IDs related to light reactions, chloroplast structure, and carbon fixation (Supplementary Dataset SD4) were used to identify a core set of 115 photosynthesis-associated orthogroups.

The index was calculated by determining the percentage of the total normalized transcriptome represented by these photosynthesis-associated orthogroups. Raw counts were normalized using size factors estimated with the estimateSizeFactors function (v1.46; Love et al. 2014). For each library, the sum of normalized counts for the photosynthesis orthogroups was divided by the total sum of normalized counts across all libraries. To ensure cross-species comparability and robustness to outliers, this percentage was scaled to the 0–1 range using the 5th and 95th percentiles of the distribution across all samples. Values below the 5th percentile were clipped to 0, and values above the 95th percentile were clipped to 1.

### Amino acid quantification and analysis

Free amino acid analysis was performed as described in (Ansaf et al. 2025). Briefly, ∼2 to 4 mg tissue was extracted with Milli-Q purified water containing 13 internal standards and analyzed using an ultra-performance liquid chromatography-tandem mass spectrometer (UPLC-MS/MS) instrument (Waters Corporation, Milford, MA). Serial dilutions of an amino acid standard mixture were analyzed alongside the samples (in duplicate) for accurate identification and quantification. Separation was performed onÅ a Kinetex LC column (2.6 µm, C18, 100 A°, 100 × 21 mm; Phenomenex, Torrance, CA) maintained at 30 °C. The injection volume was set to 10 µl, and the flow rate was set to 0.3 ml/min. The mobile phase “A” consisted of 1 mM of the ion-pairing agent perfluoroheptanoic acid, while acetonitrile served as the mobile phase “B”. The flow gradient was set as follows for B: 98% at 0 min; 80% at 0.1 min; 60% at 2.3-3.6 min; 98% at 4.0-5.98 min. MS electrospray ionization (ESI) in positive ion mode and multiple reaction monitoring (MRM) transitions for each compound were used to acquire mass spectra. Flow gas and desolvation were set to 150 and 500 l/ h, respectively. Desolvation temperature was set to 350 °C, and the collision gas flow was set to 0.15 mL/min. After run completion, the data were retrieved and analyzed using the MassLynx data analysis software (TargetLynx XS, Waters, Inc.), exported, and back-calculated to total volume and sample weight to obtain the final amounts in nmol/mg tissue. Data is available in Supplementary Dataset SD14.

To evaluate the relationship between nitrogen remobilization and leaf developmental state, Pearson correlation analysis was performed between the *ps_index* and total free amino acid concentrations in *Tripsacum dactyloides* in R. For individual amino acid correlations, concentrations μg/mg were log_2_ (x + 1) transformed before analysis. To analyze the temporal alignment between free amino acids and proteases, the expression of senescence-associated proteases (Zhang et al. 2014), *SAG12* (OG0008818), and *SAG15* (OG0012961) was extracted from the VST-normalized expression matrix. Marker expression values were z-score transformed and overlaid with total free amino acid concentrations across the 14-week sampling period.

### Biomass and nitrogen determination

Plants were excavated from field plots and thoroughly washed with potable water to remove soil and exogenous debris from underground organs. Individuals were partitioned into aboveground and underground tissues at the soil-surface interface (Supplementary Figure 7A). Tissues were dried to a constant weight at 55°C for 7 days, and dry biomass was recorded (Supplementary dataset SD15). For elemental analysis, dried tissues were subjected to a two-stage homogenization process: a preliminary coarse grind using a commercial blender (EZ600; Blendtec), followed by fine grinding in an analytical mill (A11 Basic; IKA Works) and passage through a 2-mm sieve. Representative 5-g aliquots were submitted to Midwest Laboratories (Omaha, NE, USA) for total nitrogen determination via the Dumas combustion method.

To evaluate nitrogen remobilization and allocation across species, we calculated nitrogen allocation ratios using the total nitrogen content (g) of aboveground organs (leaves, stalks, and flowers) and underground organs (roots and rhizomes). These tissues were harvested from the same individual plants used for transcriptomic profiling and photosynthetic index estimation, to ensure alignment between molecular and physiological datasets. Total N was calculated as the product of nitrogen % by dry biomass. For Figure 2G, Nitrogen allocation to aboveground tissues was expressed as the ratio of total aboveground N to total underground N. To establish the link between leaf developmental state and nitrogen recycling in perennial species, the correlation between the photosynthetic index and the nitrogen allocation to aboveground tissues was assessed using Spearman’s rank correlation coefficient (ρ). For Figure 3A, nitrogen allocation to underground tissues was expressed as the ratio of total underground N to total aboveground N. Temporal shifts in nitrogen partitioning were assessed by comparing these ratios between early (week 7) and late (week 14) sampling periods to identify accessions exhibiting a statistically significant increase in nitrogen allocation to underground tissues. All associated biomass and nitrogen data, along with corresponding RNA-seq library identifiers, are provided in Supplemental Dataset SD15.

### Validation of the Photosynthetic Index using an independent maize dataset

We applied our photosynthetic indexing methodology to an independent dataset characterizing a developmental gradient in maize (*Zea mays*) leaves (Wang et al. 2014). Gene-level counts across 15 sections of the leaf blade (from base to tip) were processed to calculate the *ps_index* using the same transcriptomic proportioning and robust scaling methodology described above, utilizing a core set of maize-specific photosynthesis-associated genes. We performed Pearson correlation analysis to evaluate the relationship between the *ps_index* and several physiological and metabolic markers measured across the same developmental gradient, including total protein, free amino acids, RuBisCO activity, chlorophyll a, glucose, and starch.

### Gene co-expression network analysis

Weighted gene co-expression network analysis (WGCNA) was performed using the WGCNA R package (Langfelder and Horvath 2008). All analyses were performed using R version 4.4.1, WGCNA version 1.73, and DESeq2 version 1.46.0. To identify regulatory networks associated with active nitrogen recycling, we focused the analysis on a cohort of “remobilizer” species and accessions identified through physiological nitrogen measurements (*A. gerardi, P. virgatum, S. nutans*, and *T. dactyloides* accession ‘Pete’). Networks were constructed independently for leaf (324 libraries) and underground (464 libraries; rhizome and root) tissues.

Input data for the leaf network consisted of VST-normalized counts for the top 3,000 most variable orthogroups. Following the removal of orthogroups with zero Median Absolute Deviation (MAD) across samples, 2,899 orthogroups were used for network construction. For the underground network, 3,000 variable orthogroups were selected, with 2,987 remaining after MAD filtering. Signed networks were constructed using the biweight midcorrelation (bicor) function with maxPOutliers = 0.1 to enhance robustness against outliers. The soft-threshold power (β) was selected based on the scale-free topology model fit (leaf β = 9; underground β = 5). Modules were detected using the blockwiseModules function with a minimum module size of 50 and a mergeCutHeight of 0.40.

Module eigengenes (MEs) were calculated as the first principal component of each module. To classify modules as sink- or source-associated, we calculated the Spearman correlation coefficient (ρ) between MEs and the Photosynthetic Index (PI). Functional importance within modules was assessed by ranking orthogroups according to their eigengene-based connectivity (kME), defined as the correlation between an orthogroup’s expression profile and its module eigengene. To visualize the global network structure, we utilized the TOM-UMAP embedding method (Morabito et al. 2023), which projects the features of the topological overlap matrix (TOM) into a two-dimensional space. For biological interpretation, modules were subjected to GO enrichment analysis as described above. All module assignments and hub rankings are available in Supplementary Dataset SD5 (leaf) and SD8 (underground). The script for the WGCNA analysis is available on the paper’s GitHub repository.

To evaluate the conservation of the co-expression modules across species, tissues, and accessions, we performed module preservation analysis using the modulePreservation function in the WGCNA package. For each target accession or tissue, the co-expression modules from the reference “remobilizer” networks (leaf or underground) were projected into the corresponding test dataset. Before analysis, test datasets were filtered to retain orthogroups with a non-missing fraction > 0.5 (*minOKFrac*).

Preservation was assessed using 100 permutations per test. We calculated the Z_summary_ statistic, which integrates multiple measures of module density and connectivity to provide a global estimate of preservation (Langfelder et al. 2011). Modules were considered strongly preserved if Z_summary_ > 10, moderately preserved if *2 <* Z_summary_ *< 10*, and not preserved if Z_summary_ < 2 (Langfelder et al. 2011). To facilitate crossspecies comparisons, values for annual species (*Zea mays* and *Sorghum bicolor*) were plotted against module size, and results were faceted by species or tissue type using custom R scripts. To evaluate the stability of intramodular topology across different life histories, we performed cross-species connectivity analysis. We calculated *kME* values for all orthogroups in the tested modules independently for annual (*Zea mays, Sorghum bicolor*) and perennial (*A. gerardi, P. virgatum, S. nutans*, and *T. dactyloides*) accessions. Preservation of the network’s internal structure was assessed by calculating the Pearson correlation between *kME* values for annual and perennial species. All preservation summary statistics are provided in Supplementary Dataset SD16-17.

## Data availability

The Expression Atlas can be accessed and explored interactively on an online website: <https://maizegenetics.github.io/atlas-of-grass-senescence-dashboard/>. The raw sequencing read data in this study have been submitted to the NCBI BioProject database (https://www.ncbi.nlm.nih.gov/bioproject/) under accession number PRJNA1458764. All custom scripts mentioned throughout methods and used to process sequencing data are available in the GitHub repository: https://github.com/jonathan-or/grass-senescence-atlas.

## Author contributions

EO, ESB, MCR, and JOOR contributed to the project’s conception and experimental design. Fieldwork and sampling were conducted by EO, JOOR, SKH, NL, TL, MCR, and ESB, with field management and logistical support from EO, NL, and MCR. JOOR performed the primary data and bioinformatic analyses with support from SKH; specialized bioinformatic assistance was provided by MCS for orthogroup inference and multispecies integration, and by JZ for count aggregation and multispecies handling scripts. Laboratory analyses were carried out by TL (high-throughput nucleic acid extraction) and AY and RA (amino acid quantification), while ESB and MCR provided reagents and analytical tools. JOOR wrote the initial draft of the manuscript, and all authors edited and approved the final version.

## Acknowledgments and funding

The authors would like to thank Sakiko Okumoto and Wolfgang Busch for initial discussions of the project. The authors would also like to thank Zachary Miller for providing technical assistance with Helixer genome annotation. This project was supported in part by Foundation for Food & Agriculture Research grants 22-000283 and 25-001855, which also included support from The Grantham Foundation for the Protection of the Environment. The project was also supported by the USDA-ARS as part of CERCA - Circular Economy that Reimagines Corn Agriculture (8062-21000-052-000-D). E.O. was supported by the United States-Israel Binational Agricultural Research and Development Fund through the Vaadia-BARD Postdoctoral Fellowship Award No. FI-628-2022.

