## Supplementary Figures for "A transcriptomic atlas of grass senescence reveals divergent underground sink networks limit nitrogen recycling in annuals"

Ojeda-Rivera et al.

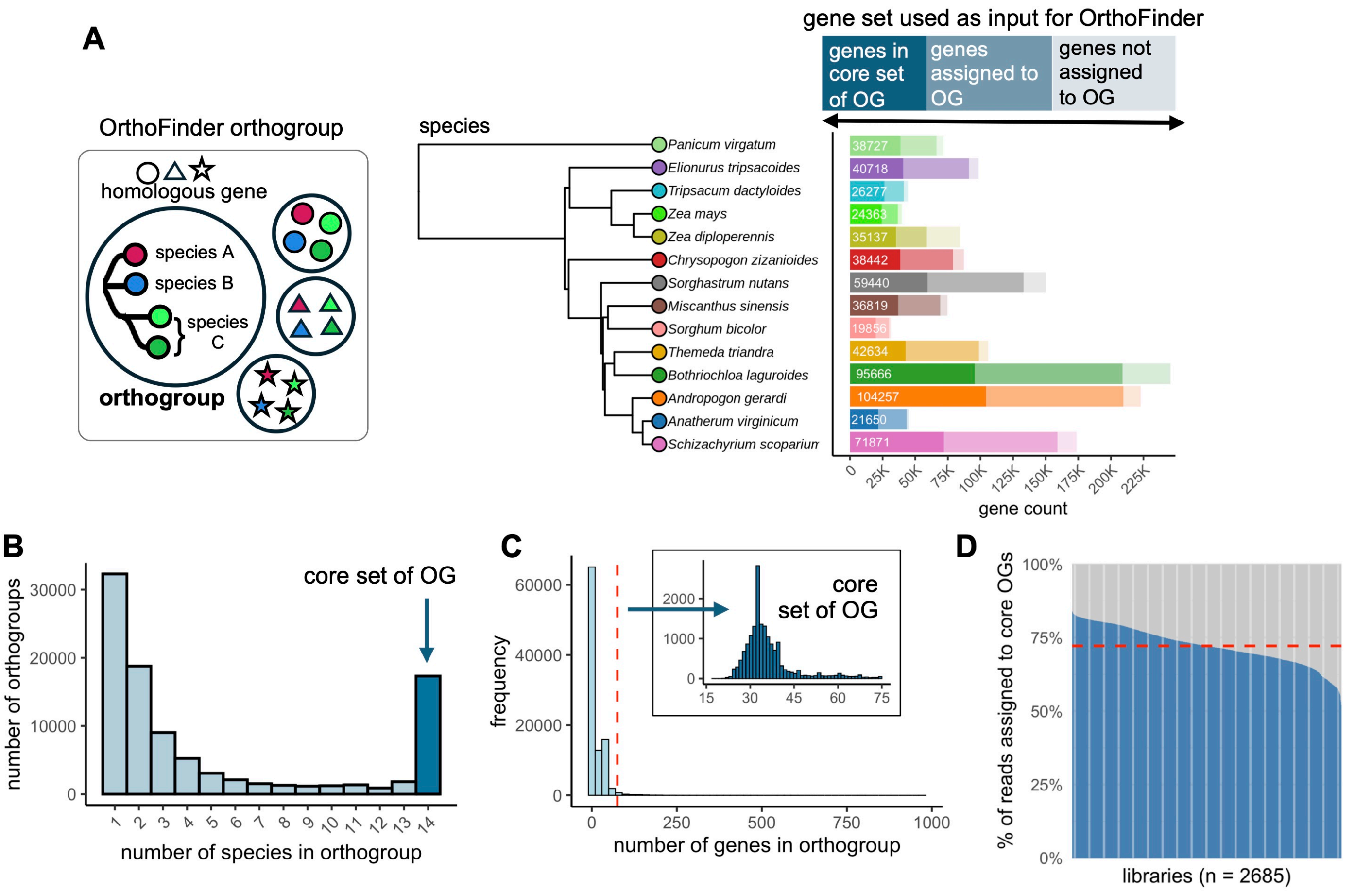


**Supplementary Figure 1. Definition of a core set of orthogroups for comparative transcriptomic analyses.**
**(A)** Summary of orthogroup assignment across 14 species inferred using OrthoFinder, showing the proportion of annotated genes not assigned to orthogroups, assigned to orthogroups, and retained in the core set. A schematic with a generic example of how OrthoFinder
**(B)** Histogram of species representation (x-axis) and number of orthogroups (y-axis), highlighting orthogroups with at least one gene member present in all species (this was one of two criteria used to define the core set).
**(C)** Histogram of gene family size (number of genes per orthogroup) for the full orthogroup set and the core set. A maximum threshold of 75 genes per orthogroup was used as the second criterion for defining the core set.
**(D)** Proportion of total read counts across all transcriptomic libraries captured by the core orthogroup set.

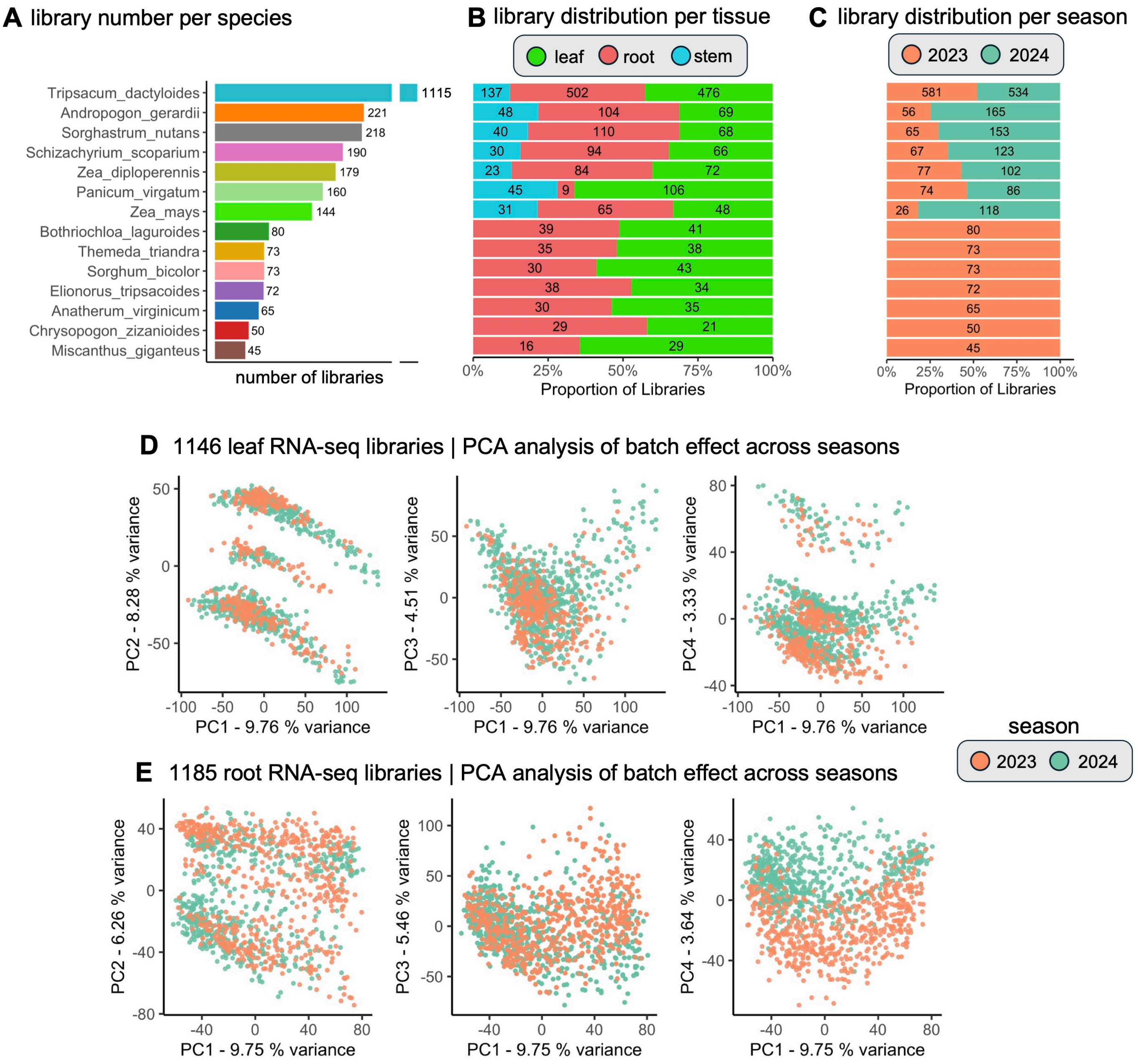


**Supplementary Figure 2. Distribution per species, tissues, and batch effect assessment per season of RNA-seq libraries.**
(A) Total number of RNA-seq libraries generated across 14 grass species.
(B) Breakdown of libraries by tissue type (leaf, root, and stem).
(C) Number of RNA-seq samples per growing season for years 2023 and 2024.
Bar width represents the relative proportion of libraries, while absolute counts are displayed within each bar segment in panels B and C. The y-axis (species) is shared across panels A, B, and C.
**(D-E)** Principal Component Analysis (PCA) of seasonal batch effects in leaf libraries (D) and root libraries (E), colored by growing season (2023 vs. 2024). In leaf libraries, there is no observable batch effect across PC1-4. For roots, season-associated differences are primarily captured by PC4, which explains 3.64% of the total variance, indicating a limited batch effect relative to overall transcriptional variation.

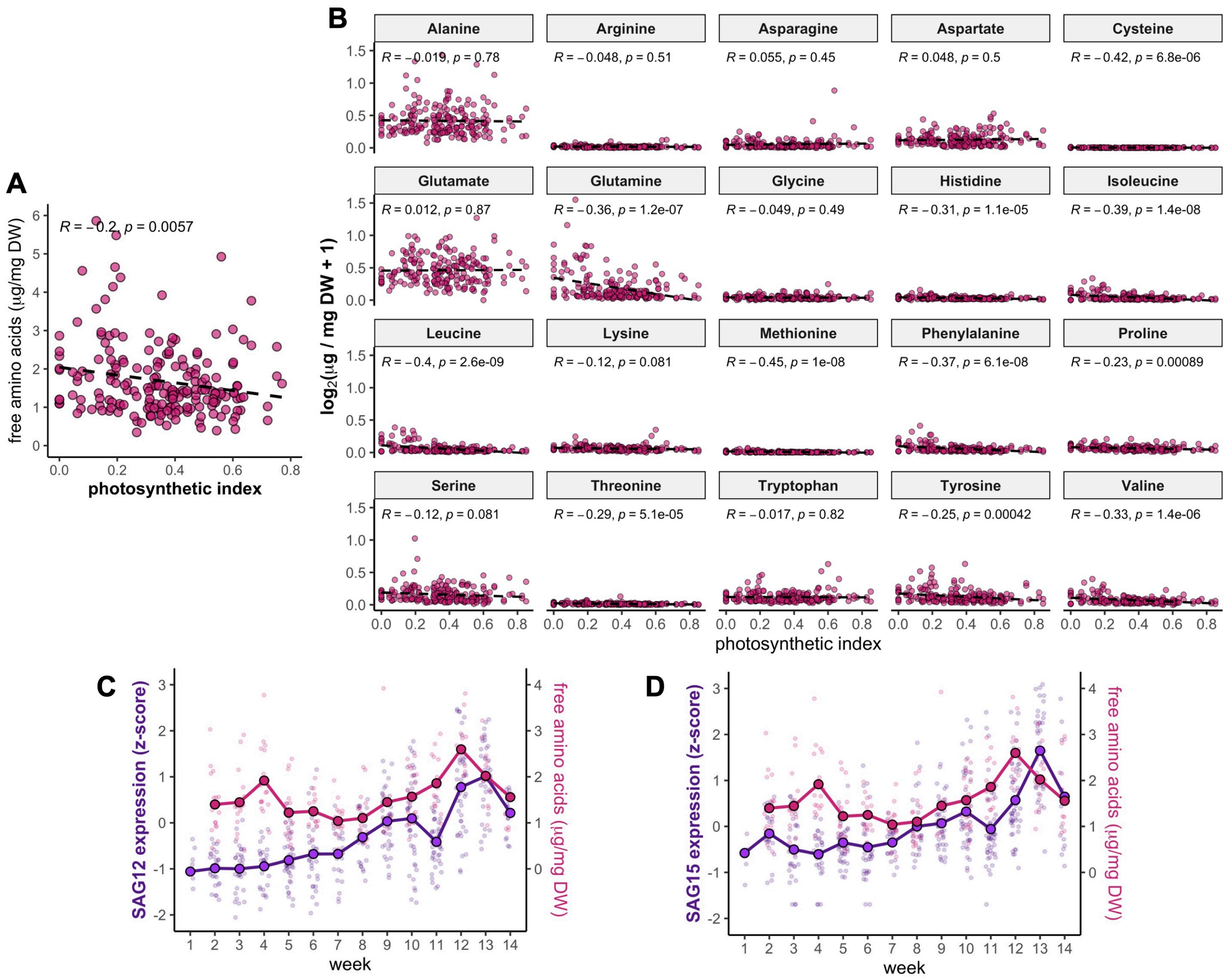


**Supplementary Figure 3. Free amino acids correlate negatively with the photosynthetic index and align with senescence protease expression in the leaves of *Tripsacum dacyloides*.**
(A) Correlation between photosynthetic index and total free amino acids in sample leaves of perennial T. dactyloides (Pearson's r is shown).
(B) Individual correlations for 20 amino acids vs. photosynthetic index. Amino acid concentrations are $\log_2$(concentration + 1) transformed.
**(C-D)** Temporal expression of orthogropus containing *SAG12* (C) and *SAG15* (D) protease-encoding genes (purple) overlaid with total free amino acid concentration (magenta) across 14 sampling weeks. Orthogroup IDs are OG0008818 (*SAG12*) and OG0012961 (*SAG15*).

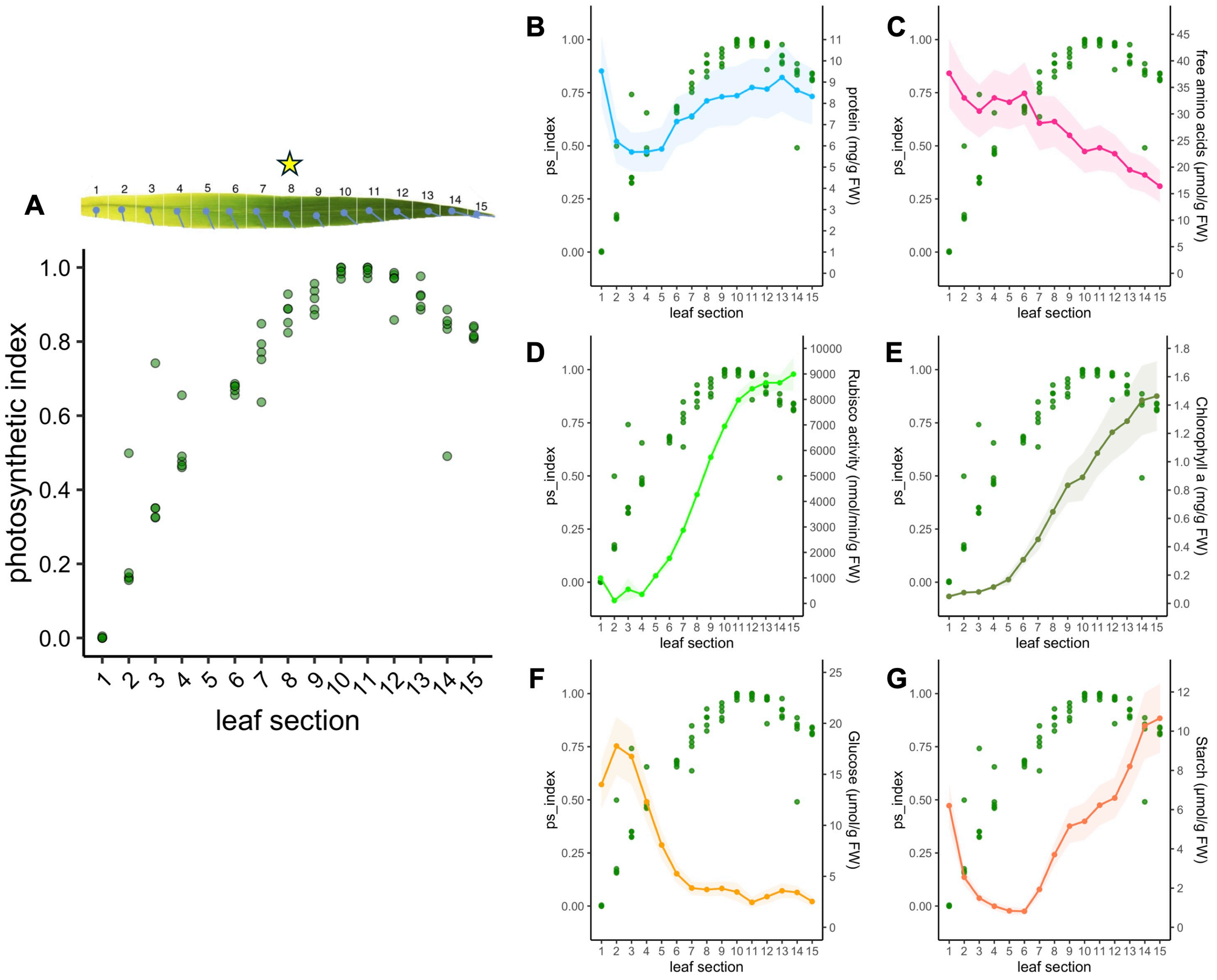


| H | protein | free amino acids | RuBisCo activity | chlorophyll a | starch | glucose |
|---|---|---|---|---|---|---|
| | r = 0.36 | r = -0.75 | r = 0.79 | r = 0.76 | r = 0.34 | r = -0.89 |

**Supplementary Figure 4. Validation of the photosynthetic index across a maize leaf developmental gradient.**
(A) Photosynthetic index values calculated across 15 sections of a maize leaf, from the base (section 1; youngest to the tip (section 15; oldest). Data were sourced from (Wang et al. 2014). The star indicates the leaf section that was sampled (for this study). **Reference**: Wang L, Czedik-Eysenberg A, Mertz RA, Si Y, Tohge T, Nunes-Nesi A, Arrivault S, Dedow LK, Bryant DW, Zhou W, et al. Comparative analyses of C  and C  photosynthesis in developing leaves of maize and rice. Nat Biotechnol. 2014:32(11):1158--1165. https://doi.org/10.1038/nbt.3019
**(B-G)** Relationship between the photosynthetic index (green points) and physiological/metabolic markers (colored lines) across the leaf gradient: total protein (B), free amino acids (C), RuBisCO activity (D), chlorophyll a (E), glucose (F), and starch (G). Lines represent mean values with shaded areas indicating standard error.
(H) Pearson correlation coefficients (r) between the photosynthetic index and the corresponding physiological or metabolic markers are shown in panels B--G.

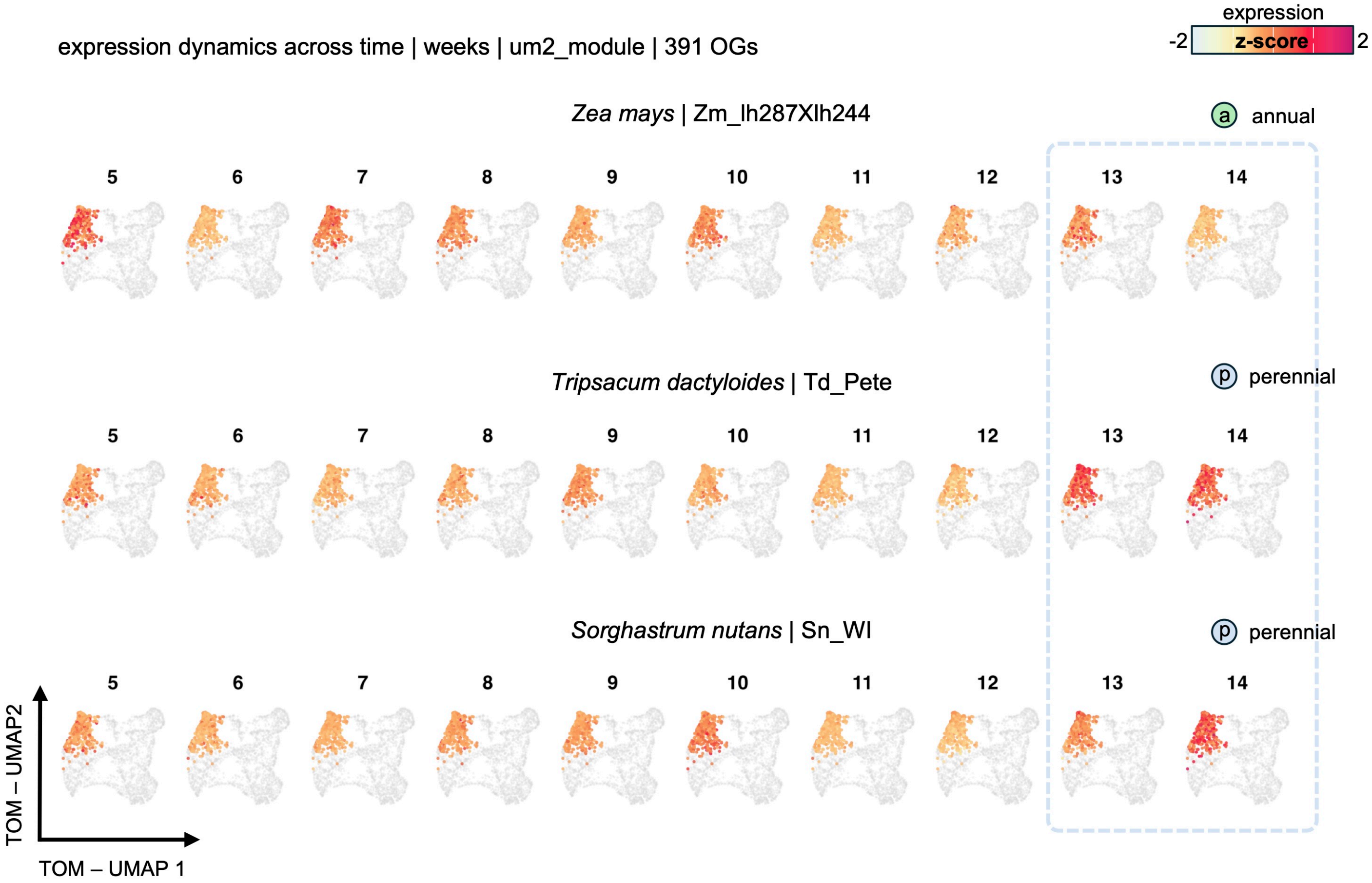


**Supplementary Figure 5. Divergent expression of the underground seed-like program in annual and perennial grasses.**
Temporal expression dynamics (weeks 5--14) of the um2_blue sub-network in Zea mays (annual), Tripsacum dactyloides(perennial), and Sorghastrum nutans (perennial), mapped onto the TOM-UMAP topology established in Figure 5E. Nodes represent orthogroups. Node colors indicate the scaled expression (z-score) across the network. The dashed blue box highlights the divergence in late-season expression (weeks 13--14), where the desiccation/dormancy program is strongly induced in perennials but absent or reduced in the annual reference.

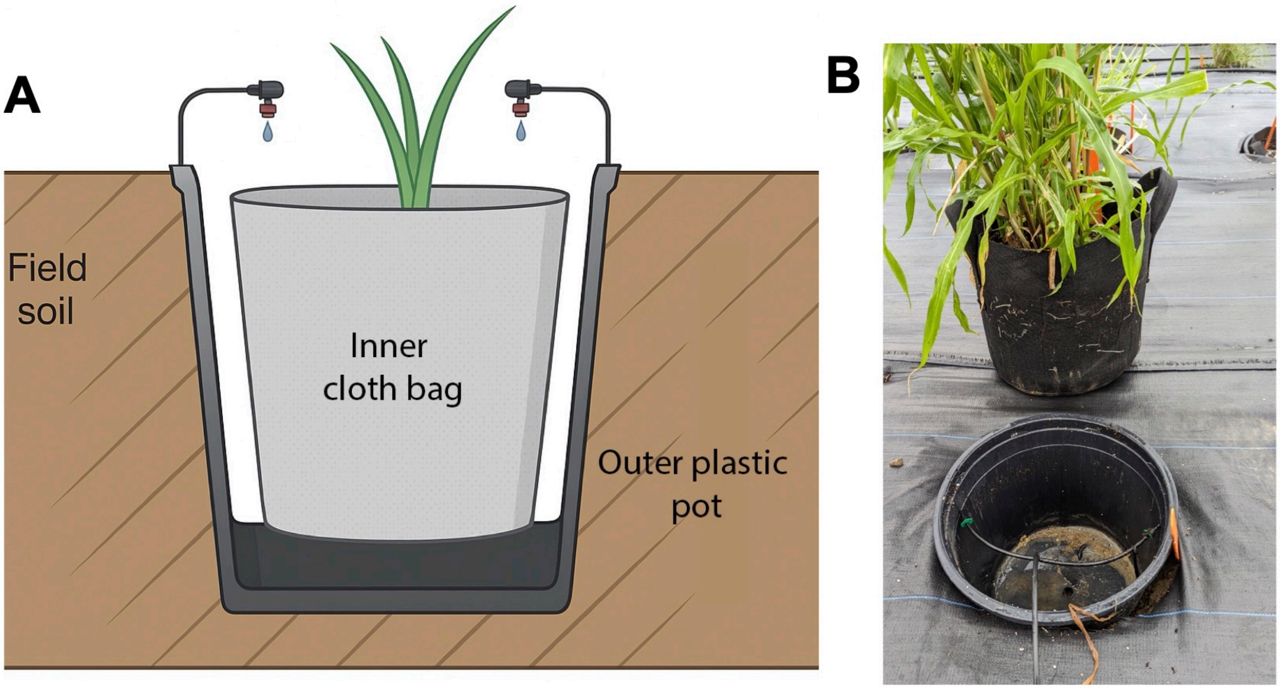


**Supplementary Figure 6. Pot-in-pot system.**
(A) Schematic diagram of the pot-in-pot system employed to grow grass accessions.
(B) Photograph of a grass accession grown in the pot-in-pot field.

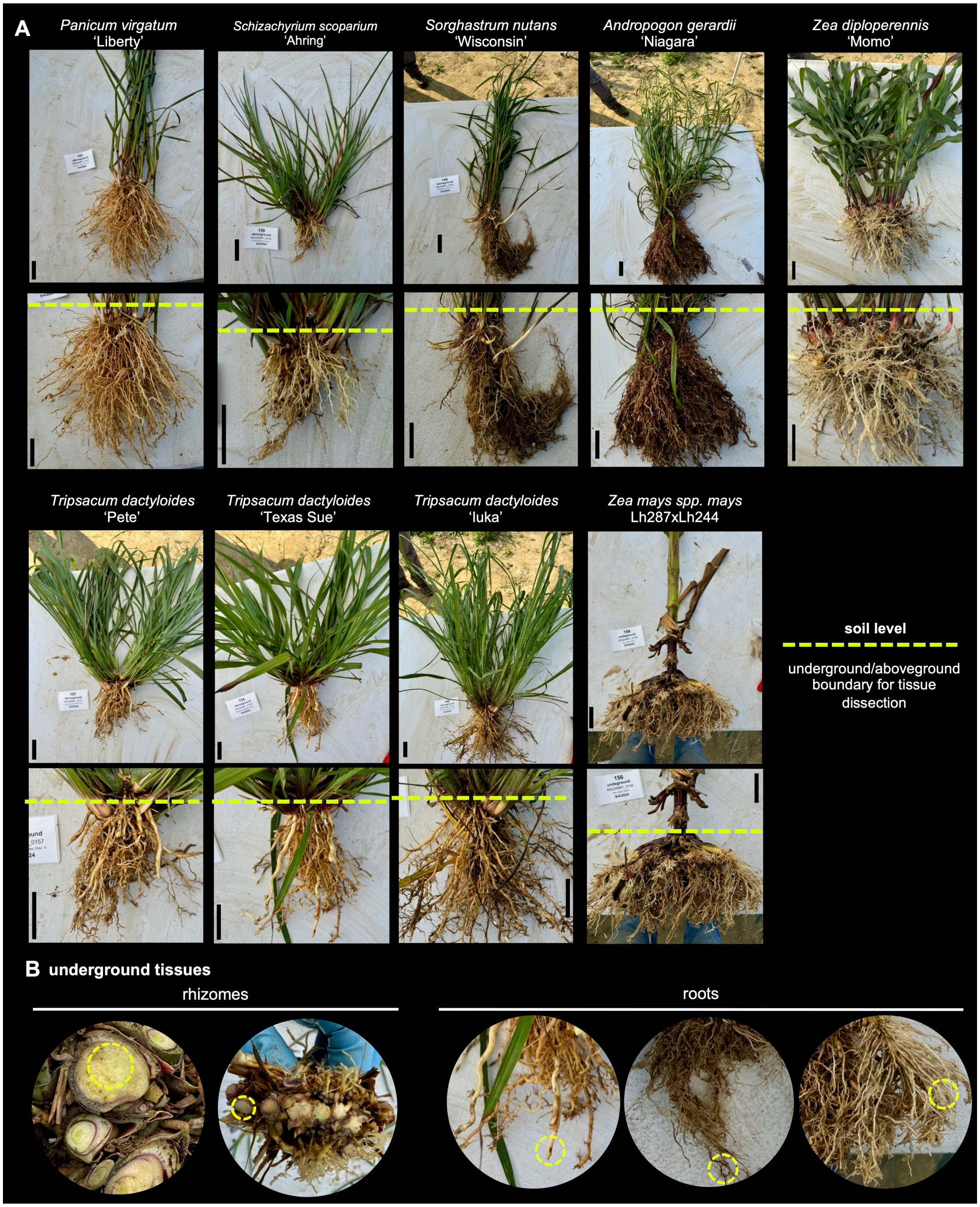


**Supplementary Figure 7. Field sampling strategy.**
(A) Grass accessions excavated from field plots, washed to expose underground organs, and dissected into aboveground and underground tissues for RNA sequencing and/or nitrogen quantification. Scale bar = 5 cm.
(B) Representative underground tissues and dissected sections used for downstream analyses. Circle diameter = 1 cm.

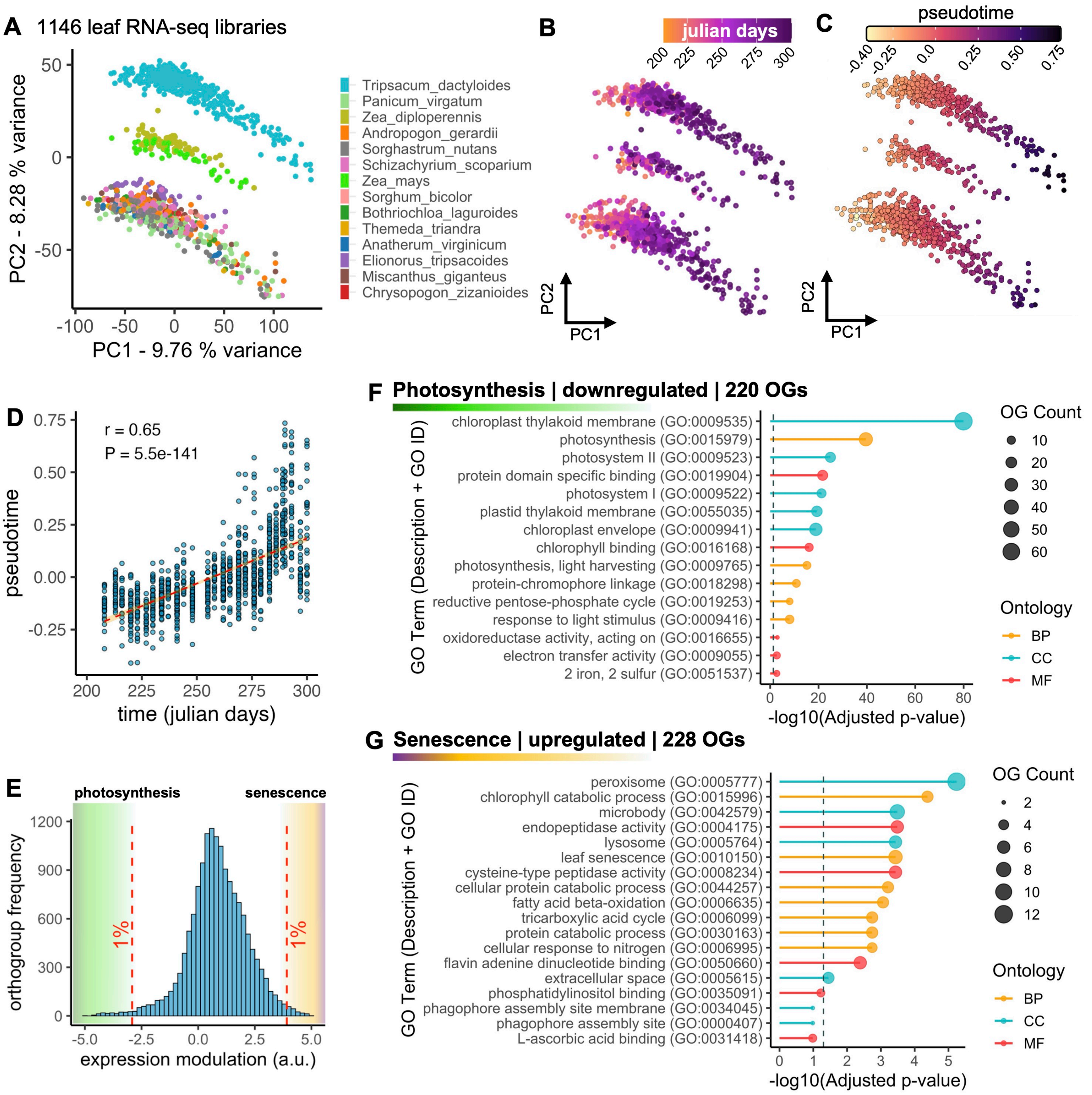


**Supplementary Figure 8. A pseudotime captured the leaf developmental transition from photosynthesis to senescence across 14 grass species.**

(A) Principal component analysis (PCA) of leaf RNA-seq libraries colored by species

(B) PCA of leaf libraries colored by sampling time.

(C) PCA of leaf libraries colored by pseudotime values.

(D) Pseudotime (y-axis) and time (Julian days) values of leaf libraries.

(E) Histogram showing the frequency distribution of orthogroup modulation values along the pseudotime trajectory (arbitrary units). Top 1% of orthogroups are highlighted.

(F) Gene Ontology (GO) enrichment analysis of orthogroups downregulated along the pseudotime trajectory.

(G) GO enrichment analysis of orthogroups upregulated along the pseudotime trajectory.
